# Cell-CLIP: Multi-algorithm causal discovery of directed cell program interaction networks from single-cell transcriptomics

**DOI:** 10.64898/2025.12.01.691666

**Authors:** Mohammed Khodor Firas Al-Tal, Xiong Liu, Joseph Zhou

**Affiliations:** Oncology Data Science, Novartis Biomedical Research, Cambridge, MA, USA; RX AICS NLP & Gen AI, Novartis Biomedical Research, Cambridge, MA, USA; Suffolk University, Boston, MA, USA

**Author notes:** **Correspondence: Joseph Zhou**, Oncology Data Science, Novartis Biomedical Research, Cambridge, MA 02139, USA.

## Abstract

Cell-cell communication networks govern tissue function in health and disease, yet existing computational tools rely on static ligand–receptor databases or correlation-based analyses that cannot distinguish direct causal relationships from confounded or indirect associations. We introduce Cell-CLIP (Cellular Causal Learning and Inference Pipeline), a modular framework that combines (i) mask-aware patient-level JASMINE activity scoring with low-percentile imputation in raw activity space, (ii) five complementary causal-discovery algorithms — PC, GES, FCI, DirectLiNGAM, and GRaSP — each fed an algorithm-matched per-column transform of the same masked JASMINE matrix, (iii) direction-preserving bootstrap aggregation with a weighted two-phase consensus, (iv) Joint Causal Inference (JCI) for principled pooling of patient cohorts via context indicators, and (v) a residual-independence (RESIT) direction audit, all evaluated under a systematic three-by-three-by-three hyperparameter sweep. Applied to a 611-patient multi-phenotype lung atlas spanning normal tissue, COVID-19 pneumonia, and lung cancer, the cross-cohort joint-causal-inference graph constructed by stratifying patients into normal, COVID-19, and tumour contexts before pooling recovers 100% of evaluable curated ground-truth edges (8/8; Wilson 95% CI 0.68–1.00) with 75.0% direction accuracy (6/8; 95% CI 0.41–0.95) and zero forbidden-orientation violations (0/8; 95% CI 0.00–0.32) — the only configuration in a comprehensive ablation series spanning 7 cohort/pool configurations, 27 hyperparameter settings, and 3 consensus types to simultaneously reach all three optimal Pareto corners. The single-cohort universal cross-phenotype graph (12 cross-phenotype programs over the full 611-patient atlas) similarly achieves 100% evaluable recall (8/8; 95% CI 0.68–1.00) with 62.5% direction accuracy (5/8; 95% CI 0.31–0.86) and zero forbidden violations (0/3; 95% CI 0.00–0.56) under the same systematic hyperparameter sweep with low-percentile imputation, exceeding an earlier zero-imputation baseline configuration of the pipeline on every reported metric (Table 6, Table 8); given small ground-truth denominators (*n*_eval = 8), these point-estimate improvements lie within the Wilson 95% intervals of the baseline and should be read as direction-of-effect rather than as confirmatory comparisons. We further show that single-cohort causal inference on the disease-extended pooled cohorts (which apply the COVID- and cancer-extended program sets to all 611 patients) recovers strong skeleton recall (35.0%–55.2% evaluable) but suffers a structural direction collapse (down to 14.3% direction accuracy and 60% forbidden rate at union) caused by within-cohort averaging over heterogeneous normal-plus-disease patient populations; the patient-stratified joint-causal-inference pool dissolves this collapse without altering the underlying algorithms. Cell-CLIP provides a hypothesis-generating framework for inferring directed cell-program networks from patient-level scRNA-seq atlases, with all reported relationships requiring experimental validation due to observational data limitations.

## Introduction

Tissue function in health and disease depends critically on cell-cell communication networks^1–3^. In normal tissue, homeostasis requires coordinated interactions among immune sentinels, stromal scaffolds, and epithelial barriers^4^. In pathological states — infection, autoimmunity, cancer — aberrant cell crosstalk drives disease progression and therapeutic resistance^5^–^7^. Understanding these context-dependent networks is essential for predicting how tissues respond to perturbations and for designing interventions that modulate microenvironmental interactions rather than isolated molecular targets^8^,^9^.

Current computational tools for inferring cell-cell communication from single-cell RNA-seq data have critical limitations^10–12^. Most methods (CellChat^13^, CellPhoneDB^2^, NicheNet^3^) rely on static ligand-receptor databases, assuming expression implies functional interaction regardless of tissue context. Two structural limitations recur across ligand–receptor-pair-level cell-cell-communication tools (CellChat^13^, CellPhoneDB^2^, NicheNet^3^, and the more recent MultiNicheNet^67^, Tensor-Cell2Cell^68^, and LIANA^69^): individual ligand-receptor expression measurements in scRNA-seq are dominated by single-gene dropout noise, and ligand-receptor binding events do not, in themselves, translate to downstream cellular function. Cell-CLIP operates one functional aggregation above this layer — on rank-aggregated cell-program activity scores that represent functional modules — mitigating both issues. Intracellular gene-regulatory-network inference (e.g., SCENIC^70^, GENIE3^71^) targets gene-to-gene edges within a single cell type and is therefore complementary rather than competitive; at the patient-level cell-program activity layer that Cell-CLIP targets, existing methods are correlation- or co-expression-based (weighted gene co-expression network analysis on program scores, GSVA combined with partial correlation) and undirected, leaving directed inference at this layer essentially unaddressed. Methods that aggregate genes into pathways and compute correlation networks^14^ address single-gene noise but cannot account for confounding: shared upstream regulators create spurious correlations, while indirect associations conflate direct and mediated effects. Weighted gene co-expression network analysis^48^,^49^ identifies co-expression modules but fundamentally cannot distinguish direct from indirect relationships^50^. Marginal correlation-based methods cannot answer the directional questions essential for mechanistic interpretation: when programs A and B are co-active, does A drive B, does B drive A, or do both depend on a shared upstream regulator?

Pearl’s causal-inference framework^15–17^ provides principled tools for discovering directed dependencies from observational data. The framework represents systems as directed acyclic graphs (DAGs) where nodes are variables and edges represent direct causal effects^18^. Modern causal-discovery algorithms infer DAG structure through conditional-independence testing (constraint-based methods)^19^, Bayesian model scoring (score-based methods)^21^, latent-variable detection (FCI)^51^, exploitation of non-Gaussianity (LiNGAM)^53^, and permutation-based scoring (GRaSP)^55^. Each algorithm class makes distinct assumptions and outputs different graph types — CPDAGs, PAGs, or DAGs — providing complementary views of the underlying causal structure^23^. A central practical question for biological applications is how to combine these views, account for missing or partially observed entries in patient-level activity matrices, and generalise across cohorts that differ systematically in disease state.

We introduce Cell-CLIP, a framework that addresses these three issues jointly. First, single-cell atlases produce per-patient activity matrices that are systematically masked: a given patient may have too few cells of a given type for a given program to be scorable. Naive zero-imputation in the raw activity space introduces a point-mass that artefactually inflates partial correlations; mean-imputation in a normalised space erases the informative-missing signal that distinguishes “absent cell type → no program activity” from “present cell type → measured low activity”. Cell-CLIP introduces a mask-aware JASMINE scoring step with three imputation policies and selects a low-percentile imputation policy in raw activity space (the column 1st-percentile injected before any rank transform) that preserves both mask awareness and the absent-cell-type signal. Second, the five candidate causal-discovery algorithms differ in the data distribution they require: linear-Gaussian methods (PC, GES, FCI, GRaSP) assume standard-normal columns, while DirectLiNGAM requires non-Gaussian columns to achieve direction identifiability. Cell-CLIP applies an algorithm-matched per-column transform to the same masked, low-imputed JASMINE matrix — inverse-normal transform (INT) for the four linear-Gaussian methods and rank-to-Student-t (df = 3) quantile transform for DirectLiNGAM — so that each algorithm sees the input matched to its identifiability assumption. Third, real patient cohorts span multiple disease states; pooling them into a single matrix introduces context confounding, while learning per-cohort graphs and post-hoc intersecting them is statistically incorrect for a pooled inference. Cell-CLIP integrates Joint Causal Inference^26^, which augments the data matrix with one-hot context indicators and forbids edges into context indicators as background knowledge, yielding a single principled multi-cohort DAG.

Applied to a 611-patient lung atlas spanning normal tissue, COVID-19 pneumonia, and lung cancer, Cell-CLIP produces a cross-context causal graph from the patient-stratified joint-causal-inference pool that recovers all evaluable curated ground-truth edges with high direction accuracy and zero forbidden-orientation violations, while a parallel single-cohort universal cross-phenotype graph reaches the same recall corner. We further use Cell-CLIP to characterise a structural failure mode of single-cohort causal inference on disease-extended pooled cohorts — within-cohort averaging over heterogeneous normal-plus-disease patient populations destroys directional signal even when skeleton recovery improves — and show empirically that joint-causal-inference pooling on patient-stratified cohorts dissolves this failure. The framework, ablation series, and curated ground truth are released to enable systematic causal discovery on future single-cell atlases.

## Methods

### Data and preprocessing

We analysed a multi-phenotype lung atlas comprising 611 patients across three conditions: normal tissue (n = 370), COVID-19 pneumonia (n = 88), and lung cancer (n = 153). All three patient cohorts were sourced from the CZ CELLxGENE Discover platform^24^. Single-cell RNA-seq data were preprocessed using standard quality control including doublet removal^26^ and library-size normalisation^27^ (total counts per cell rescaled to 10^4^, followed by a log(1 + x) transformation). We curated cell programs representing biologically coherent gene sets spanning immune effector functions, immune regulation, epithelial responses, malignant programs, and stromal programs^28^,^29^, with each program defined by marker genes derived from literature and canonical pathway databases^30–32^. Cell-type labels were normalised across studies (for example, pneumocyte → epithelial; alveolar macrophage / monocyte → macrophage) so that programs scored by cell-type-aware aggregation could be compared across patients with heterogeneous annotations.

### Mask-aware JASMINE scoring

For each (patient, cell-type, program) triple the JASMINE^47^ score is computed by rank-aggregating normalised expression of the program’s gene set within the eligible cells. Specifically, for each eligible cell JASMINE ranks every gene by its normalised expression across all cells of the target type in that patient, and the per-cell program score is the mean rank of the program’s marker genes (normalised to [0, 1]); the per-patient activity score is then the arithmetic mean of these per-cell scores across all eligible cells of the target type. We require ≥ 10 eligible cells of the matching cell type for the score to be reported; entries failing this threshold are recorded as missing values and tracked in a Boolean mask matrix. Three scoring variants were implemented: (i) **exact cell-type matching**, requiring the program’s target cell type to match exactly (the variant used throughout the headline analyses); (ii) **lineage-broadened cell-type matching**, which broadens the cell-type match to a small lineage family of related cell types (for example, all macrophage subtypes scored together for macrophage-specific programs); and (iii) **cell-type-agnostic aggregation**, which averages all cells of a patient regardless of cell type — a form of pseudobulk aggregation retained as a sensitivity diagnostic only. We compute one (patient × feature) matrix and one mask per phenotype, then merge them across phenotypes to produce a 611 × 40 patient × cell-type-program activity matrix. All downstream cohort-filtered and patient-stratified matrices analysed in this paper are derived from this merged matrix with their masks carried through. Program activity scores reflect cross-patient variation in gene-set expression across the patient population; they do not represent temporal dynamics or within-patient longitudinal changes, and causal edges in the learned graphs should be interpreted as statistical dependencies across patients rather than time-ordered mechanisms.

### Low-percentile imputation and per-algorithm transform dispatch

Causal-discovery algorithms cannot consume missing values directly. Importantly, most missing values in Cell-CLIP reflect a biological reality — the target cell type is absent or too sparse in a given patient — rather than technical measurement dropout; a patient with too few macrophages cannot have a macrophage-program score regardless of sequencing depth. Two imputation policies were compared. Median imputation on the standard-normal scale (post-INT) places masked entries at the column mean and erases the “absent cell type → no program activity” signal. Low-percentile imputation injects a low constant (the column 1st percentile in raw JASMINE space) **before** any rank transform, so that masked entries are ranked at the bottom of each column together with genuinely low-activity patients but separated from intermediate measured values. Empirically, low-percentile imputation strictly dominates median imputation on every reported cohort except the lung cancer patient-stratified cohort at majority consensus (the Sensitivity analysis section) and is used as the default throughout.

The same masked, low-imputed matrix is then dispatched per-algorithm into algorithm-matched column transforms. PC, GES, FCI, and GRaSP receive the column-wise inverse-normal transform (INT) — the standard-normal input that Fisher’s-Z partial correlation and BIC scoring assume. DirectLiNGAM receives a rank-to-Student-t (df = 3) quantile transform that is symmetric, mean-zero, and heavy-tailed (median kurtosis ≈ 3.8), restoring the non-Gaussian marginals that LiNGAM’s pairwise independence test requires for direction identifiability. The framework also exports the raw JASMINE and a logit-of-raw transform for sensitivity analysis but these are not used for the headline runs.

### Multi-algorithm causal discovery

Cell-CLIP applies five complementary causal-discovery algorithms, each exploiting different statistical properties:

#### PC algorithm

(constraint-based)^19^,^20^: discovers causal structure through iterative conditional-independence testing using Fisher’s z-transform of partial correlations (varying α ∈ {0.01, 0.025, 0.05} in the hyperparameter grid; maximum conditioning-set size = 3). Outputs a CPDAG representing a Markov equivalence class. A kernel-based independence test (KCI) is also wired in for non-linear dependencies but is not used in headline runs because of its prohibitive runtime (the Discussion).

#### GES algorithm

(score-based)^21,22^: searches DAG equivalence classes via greedy forward-backward phases, maximising the local Bayesian Information Criterion (BIC). Outputs a CPDAG. On the densest cohorts (the lung cancer patient-stratified cohort, the cancer-extended pooled cohort, and the patient-stratified JCI pool) GES is automatically excluded from a given run when its single-cohort runtime exceeds the available compute budget; the consensus then operates with a four-algorithm ensemble (PC, FCI, DirectLiNGAM, GRaSP). When GES is excluded, the majority consensus threshold scales accordingly: with four contributing methods, “majority” requires a weighted skeleton vote of more than two (i.e., support from at least three of four methods after weights), preserving the “more than half” semantics. The implications of this scalability constraint are discussed in the Discussion.

#### FCI algorithm

(latent-variable-aware)^51^,^52^: extends PC to account for unmeasured confounders, outputting a partial ancestral graph (PAG) that distinguishes direct edges from latent-confounded associations via bidirected and circle-endpoint markers. The full PAG markers are emitted alongside the binary adjacency, so program pairs flagged as possibly confounded can be reported separately from genuine forbidden violations.

#### DirectLiNGAM

(functional causal model)^53^,^54^: exploits non-Gaussianity in the data-generating process to identify a unique DAG (not just an equivalence class). The rank-to-Student-t (df = 3) input restores the heavy-tailed non-Gaussianity required by its pairwise independence test (the Low-percentile imputation section).

#### GRaSP

(permutation-based)^55^: combines greedy relaxation of sparsest permutations with BIC scoring (depth = 3), providing an additional score-based perspective that complements GES.

### Direction-preserving bootstrap aggregation

Each algorithm is executed over a configurable number of bootstrap resamples (we use 100, 200, or 300 resamples in the hyperparameter sweep, denoted *n*_boot). Rather than aggregating binary edge presence, Cell-CLIP separately counts directed and undirected edge occurrences across bootstraps, preserving directional information that standard bootstrap consensus would discard. Edges are retained per algorithm if their aggregate stability exceeds a bootstrap-stability threshold (we use 0.5, 0.6, or 0.7 in the hyperparameter sweep, denoted *t*_boot).

### Two-phase weighted directional consensus

Cell-CLIP constructs consensus DAGs through a two-phase procedure with explicit per-algorithm weighting:

#### Phase 1 — Skeleton detection

For each program pair (i, j), skeleton votes are tallied across all five methods with equal per-method skeleton weights (1.0 for each method in all reported runs). An edge is included if the weighted vote exceeds the consensus threshold: full agreement (= 5.0 with five methods, = 4.0 with four) for **intersection**, more than half for **majority**, and ≥ 1.0 for **union**.

#### Phase 2 — Direction determination

For edges passing the skeleton filter, directional votes are computed as weighted sums of per-method direction weights *w*_dir: fully directed edges contribute *w*_dir votes in their direction; undirected edges contribute *w*_dir/2 votes in both directions. The direction weights are 1.0 for each linear-Gaussian method (PC, GES, FCI, GRaSP) and 1.5 for DirectLiNGAM, reflecting that DirectLiNGAM is the only method in the ensemble that can break Markov-equivalence-class ties via non-Gaussianity. The 1.5× value was chosen empirically as a conservative single-step promotion (the natural alternatives are 1.0× — equal weight — and 2.0× — full priority over a single tying linear-Gaussian vote) and was used uniformly across all reported runs; no value other than 1.0× and 1.5× was tested in isolation. The 1.0× setting is included as one component of the LiNGAM-direction-downgrade hybrid sensitivity variant in the Sensitivity analysis section, which combines this direction-weight reduction (1.5× → 1.0×) with an inverse-normal-transform fallback applied to low-fill columns; this combined sensitivity variant produced no improvement on any reported cohort and a measurable regression on the lung cancer patient-stratified cohort, so the 1.5× setting is retained as the production default. A cleaner isolated comparison (1.0× direction weight without the inverse-normal-transform-fallback change) was not run within the wall-time budget of this study and is flagged as a target for future DirectLiNGAM direction-weight sensitivity work (the Discussion). The direction with more votes wins; remaining ties are annotated as bidirectional.

Three consensus DAGs are produced per (cohort, hyperparameter tag): **intersection** (highest specificity), **majority** (balanced), and **union** (highest sensitivity). The reporting consensus is selected per-cohort based on the Pareto front of validation metrics (the Pareto fronts section).

### DAG conversion and acyclicity enforcement

Consensus graphs are converted to proper DAGs via a procedure that uses LiNGAM adjacency matrices as priors to orient any remaining undirected edges. Acyclicity is enforced through depth-first cycle detection with iterative removal of the weakest back-edge by bootstrap support, guaranteeing a valid DAG output.

### Joint Causal Inference for multi-cohort pooling

To combine evidence across patient cohorts in a single principled inference, Cell-CLIP implements Joint Causal Inference (JCI)^62^, an instance of the more general causal-data-fusion programme^65^ that aims to consistently integrate observations collected under heterogeneous source populations or experimental conditions. Per-context patient × program matrices are vertically stacked and the augmented design is appended with one-hot context-indicator columns (one indicator dropped for identifiability against a chosen reference context). Background knowledge forbids edges into context indicators (i.e., program → context) and edges between context indicators (context ↔ context) for the constraint-based and score-based methods that natively accept this prior (PC, GES, FCI). DirectLiNGAM does not natively accept this prior in the implementation we use; consensus arrows from program to context that originate from DirectLiNGAM are therefore retained in the augmented adjacency and reinterpreted at the reporting stage as cohort-discriminative-program markers (the Cohort-discriminative-program arrows section) rather than as causal program-to-context effects. The five-algorithm sweep then runs on the augmented matrix; the resulting adjacency is partitioned into three submatrices: program × program (the cross-context biological graph), context-indicator × program (cohort-effect arrows from a context indicator into a program), and program × context-indicator (cohort-discriminative-program arrows). Two pools are reported: a **disease-extended cross-cohort JCI pool** that stacks the universal cross-phenotype cohort, the COVID-extended pooled cohort, and the cancer-extended pooled cohort on the 12 universal program columns; and a **patient-stratified cross-cohort JCI pool** that stacks the normal-tissue patient-stratified cohort, the COVID-19 patient-stratified cohort, and the lung cancer patient-stratified cohort on the same 12 columns, with the normal-tissue cohort designated as the reference context.

### RESIT direction audit

After consensus, each directed edge is independently audited via Regression with Subsequent Independence Test (RESIT)^63^. For each consensus edge `i → j`, RESIT fits `x_j = f(x_i) + ε_ij` and `x_i = g(x_j) + ε_ji` with gradient-boosting regressors and tests whether the residuals are independent of the regressor via an HSIC permutation test^64^. Each edge is labelled **forward** (RESIT supports the consensus direction), **reverse** (RESIT prefers the opposite orientation), **ambiguous** (HSIC tolerates either direction — the data does not identify orientation under the additive-noise model), or **weak** (no orientation has independent residuals). The audit is annotation-only: it does not override consensus; it provides reviewers and downstream users with direct visibility into how much of the directed graph is supported by an independent residual-independence check.

### Hyperparameter sweep

For each headline cohort, Cell-CLIP executes a predefined three-by-three-by-three hyperparameter grid: significance level α ∈ {0.01, 0.025, 0.05} × bootstrap-stability threshold *tboot* ∈ *{0.5, 0.6, 0.7} × number of bootstrap resamples nboot* ∈ *{100, 200, 300}, producing 27 grid points × 3 consensus rules (intersection, majority, union) per cohort. Validation metrics (the Ground-truth validation section) are computed at every grid point and the Pareto front jointly over evaluable recall, direction accuracy, and forbidden-violation rate is reported, with the headline grid point chosen as the recall-maximising configuration with the lowest forbidden-violation rate (ties broken by direction accuracy). Throughout this paper individual grid points are denoted by the compact reproducibility tag aXXbtYbnNNN*, defined fully in the caption of Table 3.

### Ground-truth validation

Biological validation was performed using a curated set of directed ground-truth edges spanning three contexts (normal, COVID-19, tumour), with each edge annotated as **textbook** or **strong-evidence**, plus a separate set of **forbidden** orientations (developmentally or biologically impossible directions, e.g., cytotoxic → priming or efferocytosis → pro-inflammatory). The ground truth was assembled from KEGG^30^, Reactome^31^, and ImmuneSigDB^32^ databases supplemented with textbook immunology relationships^38–46^,^56–61^; for the patient-stratified joint-causal-inference pool the per-context ground truths are merged. Each discovered edge is classified as: **confirmed** (correct direction matches ground truth), **reversed** (skeleton correct but direction inverted), **novel** (not in ground truth, hypothesis-generating), or **forbidden** (contradicts an entry in the forbidden-orientation set). Quantitative metrics include: **evaluable recall** (the primary metric; the fraction of ground-truth edges whose two programs both appear in the learned DAG and which are recovered in the skeleton), **direction accuracy** (the fraction of evaluable recovered edges with correct direction), **forbidden-violation rate** (the fraction of evaluable forbidden edges with at least one violation), **skeleton recall** (the absolute-recall companion that does not control for program-set coverage), and **skeleton precision**. The primary metric is evaluable recall because ground-truth coverage of context-specific programs is non-uniform across cohorts. For reproducibility, these five metrics are emitted to the per-run validation files under the column names *evaluable_recall*, *direction_accuracy*, *forbidden_violation_rate*, *skeleton_recall*, and *skeleton_precision*; in the body of the paper the reader-facing English names above are used throughout. All binomial rates (evaluable recall, direction accuracy, forbidden-violation rate, skeleton recall, skeleton precision) are reported with Wilson 95% confidence intervals computed in closed form from the per-cohort numerator (*k*) and denominator (*n*); given the small evaluable ground-truth denominators in this study (8 ≤ *n*_eval ≤ 29 across cohorts), these intervals are wide and should be read as the primary calibration of statistical confidence on every reported rate.

### Software and reproducibility

Cell-CLIP was implemented in Python 3.12. Causal-discovery algorithms were executed via the `causal-learn` package33; LiNGAM via the `lingam` package; gradient-boosting regressors for RESIT via `scikit-learn`^35^. Single-cell preprocessing utilised Scanpy^34^ and AnnData^25^. Network manipulation used NetworkX^36^. All random operations used a fixed seed (42). Code, sweep logs, and curated ground-truth files are available at the location specified in the Data Availability statement. Compute scalability — and the small number of analyses constrained by it — is discussed in the Discussion.

## Results

### Dataset, program contexts, and JCI pool design

We applied Cell-CLIP to a 611-patient lung atlas across three conditions: normal tissue (n = 370), COVID-19 pneumonia (n = 88), and lung cancer (n = 153) (Table 1). From curated cell programs spanning immune, epithelial, stromal, and malignant compartments, five single-cohort matrices and two joint-causal-inference (JCI) pools were assembled.

**Table 1.**
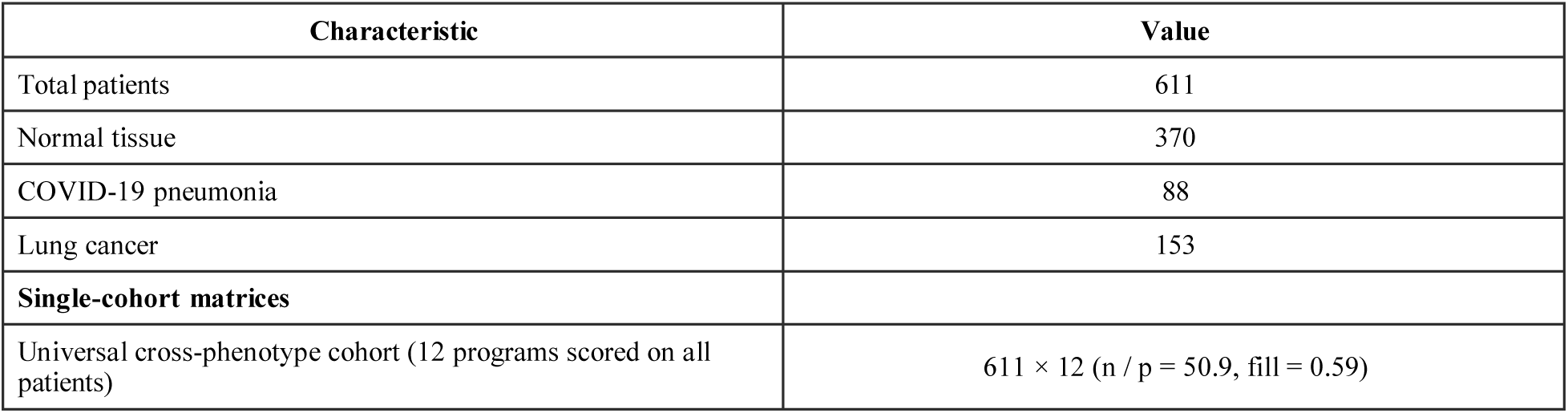
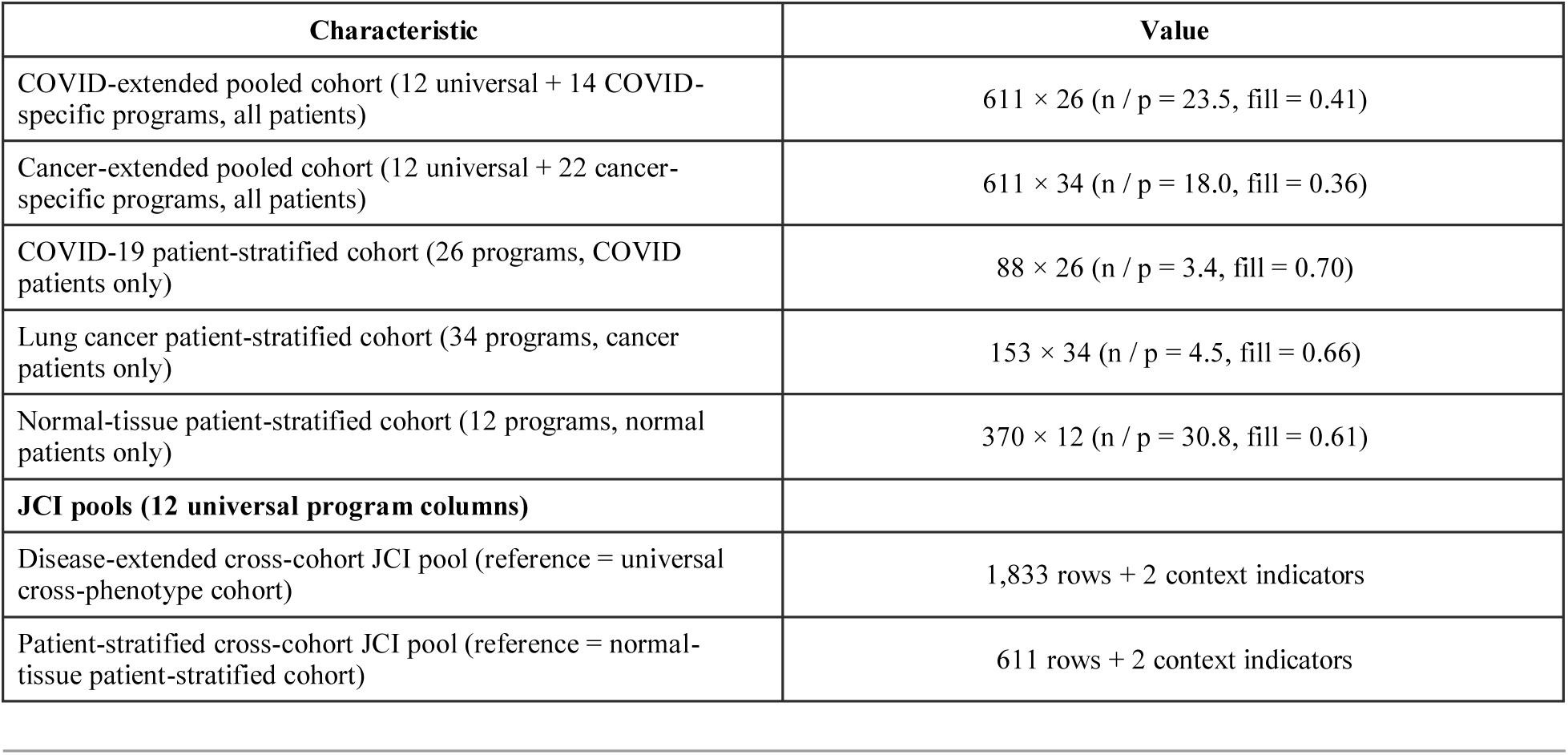
Cohort, program-context, and JCI-pool characteristics.

| Characteristic | Value |
| --- | --- |
| Total patients | 611 |
| Normal tissue | 370 |
| COVID-19 pneumonia | 88 |
| Lung cancer | 153 |
| **Single-cohort matrices** |  |
| Universal cross-phenotype cohort (12 programs scored on all patients) | 611 × 12 (n / p = 50.9, fill = 0.59) |
| COVID-extended pooled cohort (12 universal + 14 COVID-specific programs, all patients) | $611 \times 26$ (n / p = 23.5, fill = 0.41) |
| Cancer-extended pooled cohort (12 universal + 22 cancer-specific programs, all patients) | $611 \times 34$ (n / p = 18.0, fill = 0.36) |
| COVID-19 patient-stratified cohort (26 programs, COVID patients only) | $88 \times 26$ (n / p = 3.4, fill = 0.70) |
| Lung cancer patient-stratified cohort (34 programs, cancer patients only) | $153 \times 34$ (n / p = 4.5, fill = 0.66) |
| Normal-tissue patient-stratified cohort (12 programs, normal patients only) | $370 \times 12$ (n / p = 30.8, fill = 0.61) |
| **JCI pools (12 universal program columns)** |  |
| Disease-extended cross-cohort JCI pool (reference = universal cross-phenotype cohort) | 1,833 rows + 2 context indicators |
| Patient-stratified cross-cohort JCI pool (reference = normal-tissue patient-stratified cohort) | 611 rows + 2 context indicators |

#### Single-cohort matrices

The **universal cross-phenotype cohort** (611 × 12) contains the 12 cross-phenotype programs scored on every patient and is the lowest-confounding single-cohort baseline. The **COVID-extended pooled cohort** (611 × 26) and the **cancer-extended pooled cohort** (611 × 34) extend the universal cross-phenotype cohort with COVID-and cancer-specific programs respectively, scored on the full 611-patient matrix; these matrices pool all patients regardless of phenotype and are used in this paper to characterise the within-cohort mixing failure mode (the Disease-extended direction collapse section). The **COVID-19 patient-stratified cohort** (88 × 26) and the **lung cancer patient-stratified cohort** (153 × 34) restrict the same 26- and 34-program contexts to the COVID and lung cancer patients only. These matrices remove within-cohort phenotype mixing at the cost of reduced sample size. The **normal-tissue patient-stratified cohort** (370 × 12) restricts the universal program set to normal patients and is used as a reference context for joint-causal-inference pooling.

#### JCI pools

The **disease-extended cross-cohort JCI pool** stacks the universal cross-phenotype cohort, the COVID-extended pooled cohort, and the cancer-extended pooled cohort on the 12 universal program columns; the augmented design has 611 × 3 = 1,833 rows plus two context-indicator columns (one dropped against the universal cross-phenotype cohort as reference). The **patient-stratified cross-cohort JCI pool** stacks the normal-tissue, COVID-19, and lung cancer patient-stratified cohorts on the same 12 columns; the augmented design has 370 + 88 + 153 = 611 rows plus two context-indicator columns (one dropped against the normal-tissue patient-stratified cohort as reference). The two pools share an identical program set but differ in cohort composition: the disease-extended pool stacks pooled all-patient cohorts (each row averages across heterogeneous patient phenotypes), while the patient-stratified pool stacks strictly disjoint patient populations.

### Mask-aware scoring exposes informative-missing structure

The merged 611 × 40 mask-aware activity matrix is approximately 32% filled; the remaining 68% of entries are genuinely “not measured” rather than “low activity” (Table 2). Most masked entries are off-context (for example, COVID-specific programs in cancer patients) or low-cell-count (a patient lacks ≥ 10 cells of the program’s target cell type). After cohort-filtering and patient-stratification, the per-cohort fill rates rise to 0.59–0.70 (Table 2), indicating that the merged matrix’s apparent sparsity reflects cross-phenotype cell-type heterogeneity rather than a scoring deficit.

**Table 2.** Mask-aware JASMINE fill rates (exact cell-type matching).

| Matrix | Shape | Fill rate | Measured / total entries |
| --- | --- | --- | --- |
| Merged three-phenotype matrix | $611 \times 40$ | 0.323 | 7,898 / 24,440 |
| Universal cross-phenotype cohort | $611 \times 12$ | 0.587 | 4,304 / 7,332 |
| COVID-extended pooled cohort | $611 \times 26$ | 0.412 | 6,547 / 15,886 |
| Cancer-extended pooled cohort | $611 \times 34$ | 0.357 | 7,419 / 20,774 |
| COVID-19 patient-stratified cohort | $88 \times 26$ | 0.704 | 1,612 / 2,288 |
| Lung cancer patient-stratified cohort | $153 \times 34$ | 0.663 | 3,449 / 5,202 |
| Normal-tissue patient-stratified cohort | $370 \times 12$ | 0.612 | 2,717 / 4,440 |

### Multi-algorithm causal discovery: per-method edge counts

We applied Cell-CLIP across all five single-cohort matrices and the two joint-causal-inference (JCI) pools, evaluating the full hyperparameter sweep under low-percentile imputation. Per-method directed-edge counts at the headline grid points (Table 3) reveal characteristic algorithm behaviours: PC and FCI produce comparable, mid-density skeletons; GRaSP is the sparsest; GES is BIC-noisy and is automatically excluded from the densest cohorts to keep individual analyses within the available compute budget (the lung cancer patient-stratified cohort, the cancer-extended pooled cohort, and the patient-stratified JCI pool); and DirectLiNGAM consistently produces the richest skeleton, reflecting the rank-to-t (df = 3) transform restoring the non-Gaussianity its pairwise test requires. The implications of GES exclusion on the densest cohorts — and on the consequent four-method consensus threshold — are addressed as a scalability consideration in the Discussion.

**Table 3.** Per-method directed-edge counts at the headline hyperparameter grid point. Hyperparameter labels are written as `aXX_btY_bnNNN`, with `aXX` encoding the significance level α (0.01 → `a01`, 0.025 → `a025`, 0.05 → `a05`), `btY` encoding the bootstrap-stability threshold (Y ∈ {5, 6, 7} → 0.5, 0.6, 0.7), and `bnNNN` encoding the number of bootstrap resamples.

| Cohort | Headline grid point | PC | FCI | Direct LiNGAM | GRaSP | GES | Majority edges | Union edges |
| --- | --- | --- | --- | --- | --- | --- | --- | --- |
| Universal cross-phenotype cohort | $a_{025\_bt5\_bn100}$ | 18 | 17 | 29 | 15 | 27 | 19 | 28 |
| COVID-19 patient-stratified cohort | $a_{01\_bt7\_bn300}$ | 13 | 13 | 20 | 12 | 20 | 13 | 23 |
| Lung cancer patient-stratified cohort | $a_{01\_bt7\_bn300}$ | 22 | 20 | 53 | 17 | excluded (the Discussion) | 19 | 49 |
| COVID-extended pooled cohort | $a_{01\_bt7\_bn300}$ | 24 | 25 | 72 | 29 | 50 | 33 | 65 |
| Cancer-extended pooled cohort | $a_{01\_bt7\_bn300}$ | 35 | 34 | 89 | 30 | excluded (the Discussion) | 33 | 88 |
| Disease-extended cross-cohort JCI pool | $a_{025\_bt7\_bn200}$ | 25 | 25 | 32 | 21 | 30 | 24 | 30 |
| Patient-stratified cross-cohort JCI pool | $a_{025\_bt6\_bn100}$ | 20 | 20 | 38 | 15 | excluded (the Discussion) | 19 | 36 |

For the two JCI pools, per-method counts include both program × program edges and the cohort-effect edges connecting context indicators with programs; a partition between these submatrices for the headline patient-stratified pool is given in the Cohort-discriminative-program arrows section. GES exclusion on the densest cohorts is a runtime-budget consequence rather than a methodological decision; consensus on these cohorts therefore aggregates four methods (PC, FCI, DirectLiNGAM, GRaSP) and the majority threshold scales accordingly (the Multi-algorithm causal discovery section).

### Universal causal network

The single-cohort universal cross-phenotype cohort’s hyperparameter sweep produces a Pareto front with three structural regimes: a recall-maximising regime (`a025_bt5_bn100` majority, 19 edges) that recovers all eight evaluable ground-truth edges (8/8; Wilson 95% CI 0.68–1.00); a direction-maximising regime (`a01_bt6_bn100` and `a025_bt6_bn300` union, 19 / 22 edges) that achieves perfect direction accuracy (6/6; 95% CI 0.61–1.00) at recall = 0.750 (6/8; 95% CI 0.41–0.93); and an intermediate regime that interpolates the two. We adopt the recall-maximising grid point as the universal headline because its forbidden-violation rate is zero (0/3; 95% CI 0.00–0.56) and direction accuracy at full recall is 62.5% (5/8; 95% CI 0.31–0.86) — exceeding the earlier zero-imputation baseline configuration of the pipeline on every reported metric (Table 4, Table 6); given the small ground-truth denominators (*n*_eval = 8), these point-estimate gains lie within the Wilson 95% intervals of the baseline and should be read as direction-of-effect rather than as statistically significant differences.

**Table 4.** Universal headline majority DAG (a025_bt5_bn100): complete edge list.

| Source → Target | Class | Methods |
| --- | --- | --- |
| CD4 activation → germinal-centre B-cell program | confirmed (textbook) | PC, FCI, DirectLiNGAM |
| CD4 activation → CD8 cytotoxic-T program | confirmed (textbook) | PC, FCI, DirectLiNGAM, GRaSP |
| CD4 activation → NK cytotoxic program | confirmed (supportive) | multi-method |
| CD8 priming → CD4 activation | reversed (textbook ground truth: CD4 → priming) | multi-method |
| CD8 priming → CD8 cytotoxic-T program | confirmed (textbook) | PC, FCI, DirectLiNGAM, GRaSP |
| Endothelial-barrier program → germinal-centre B-cell program | novel | multi-method |
| Endothelial-barrier program → growth-factor fibroblast program | confirmed (supportive) | multi-method |
| Endothelial-barrier program → NK cytotoxic program | novel | multi-method |
| Extracellular-matrix fibroblast → endothelial-barrier program | confirmed (strong) | multi-method |
| Extracellular-matrix fibroblast → growth-factor fibroblast | confirmed (strong) | PC, DirectLiNGAM, GRaSP |
| Growth-factor fibroblast → NK cytotoxic program | novel | multi-method |
| Efferocytosis-poised macrophage → alveolar-macrophage sensing program | novel | multi-method |
| Efferocytosis-poised macrophage → CD4 activation | novel | multi-method |
| Efferocytosis-poised macrophage → macrophage recruitment program | confirmed (supportive) | multi-method |
| Pro-inflammatory macrophage → alveolar-macrophage sensing program | reversed (in reality it is bidirectional) | multi-method |
| Pro-inflammatory macrophage → germinal-centre B-cell program | novel | multi-method |
| Pro-inflammatory macrophage → efferocytosis-poised macrophage | confirmed | DirectLiNGAM |
| Pro-inflammatory macrophage → macrophage recruitment program | reversed (strong-evidence GT: recruit → pro-inflammatory) | multi-method |
| CD8 cytotoxic-T program → NK cytotoxic program | confirmed (supportive) | multi-method |

The headline universal majority graph at `a025_bt5_bn100` (Table 4) recovers five of eight evaluable curated ground-truth edges with correct orientation: CD8 priming → cytotoxic effector (textbook), CD4 activation → B-cell germinal-centre formation (textbook), pro-inflammatory macrophages → efferocytosis-poised macrophages (strong evidence), extracellular-matrix fibroblasts → growth-factor fibroblasts (strong evidence), and extracellular-matrix fibroblasts → endothelial-barrier programs (strong evidence). Three additional ground-truth edges are recovered in the skeleton with reversed orientation (CD4 activation ↔ CD8 priming; alveolar-macrophage sensing ↔ pro-inflammatory macrophages; macrophage recruitment ↔ pro-inflammatory macrophages), and one ground-truth edge is structurally non-evaluable on this cohort because one of its endpoints (the DC1 cross-presentation program) is not in the universal program set. Five additional edges have literature-supportive orientations beyond the strict curated ground-truth set (CD4 activation → CD8 cytotoxic-T program; CD4 activation → NK cytotoxic program; endothelial-barrier program → growth-factor fibroblast program; efferocytosis-poised macrophage → macrophage recruitment program; CD8 cytotoxic-T program → NK cytotoxic program). Six novel hypothesis-generating edges are produced, including a putative efferocytosis-driven CD4-activation edge that constitutes a candidate efferocytosis-feedback signal worth experimental follow-up.

#### RESIT direction audit

Independently auditing the 12 evaluable directed edges in the universal majority DAG via the residual-independence test described in the RESIT direction audit section yields 6 forward-supported edges, 1 reverse-preferred edge, and 5 ambiguous edges (the additive-noise model does not identify orientation against an HSIC permutation null). The 6 forward-supported edges include the four strongest confirmed ground-truth orientations (CD8 priming → cytotoxic effector; CD4 activation → germinal-centre B-cell formation; pro-inflammatory macrophages → efferocytosis-poised macrophages; extracellular-matrix fibroblasts → growth-factor fibroblasts) and the novel efferocytosis-driven CD4-activation hypothesis, providing residual-level support for the consensus orientations. The 5 ambiguous edges are all in the hypothesis-generating set (novel or literature-supportive beyond the curated ground truth): they represent structurally recoverable skeleton signal whose orientation cannot be determined under additive-noise residual independence and would require perturbation experiments to resolve (the Discussion).

### Cross-context causal network from the patient-stratified JCI pool — primary biological finding

All inferred edge directions in this section reflect statistical asymmetry under the distributional assumptions of the contributing algorithms (non-Gaussianity for DirectLiNGAM; Markov-equivalence-class resolution by conditional independence for PC, GES, FCI, and GRaSP) and constitute hypothesis-generating directional assessments; they are not established mechanistic causal claims and require experimental validation as stated throughout.

We computed the systematic hyperparameter sweep with low-percentile imputation on the patient-stratified cross-cohort JCI pool (the joint pool over the normal-tissue, COVID-19, and lung cancer patient-stratified cohorts on the 12 universal program columns). One configuration — α = 0.025, bootstrap threshold = 0.6, 100 bootstrap iterations on the union consensus (labelled `a025_bt6_bn100` union, with 26 program × program edges) — simultaneously recovers all eight evaluable ground-truth edges (evaluable recall = 1.000, 8/8; Wilson 95% CI 0.68–1.00), achieves 75.0% direction accuracy on those edges (6/8 oriented correctly; 95% CI 0.41–0.95), and has zero forbidden-orientation violations against eight evaluable forbidden edges (0/8; 95% CI 0.00–0.32) (Table 5). Across the entire ablation series spanning seven cohort/pool configurations, 27 hyperparameter settings, and three consensus types, this is the only configuration to simultaneously hit all three optimal Pareto corners (recall = 1.000, direction ≥ 0.750, forbidden = 0). Our primary biological finding therefore rests on this cross-context graph.

**Table 5.** Patient-stratified JCI headline DAG (a025_bt6_bn100 union): complete program × program edge list (26 edges).

| Source → Target | Class | Methods |
| --- | --- | --- |
| CD4 activation → CD8 priming | confirmed (textbook) | PC, FCI, GRaSP |
| CD8 priming → CD8 cytotoxic-T program | confirmed (textbook) | PC, FCI, DirectLiNGAM, GRaSP |
| CD4 activation → germinal-centre B-cell program | confirmed (textbook) | PC, FCI, DirectLiNGAM |
| Macrophage recruitment → pro-inflammatory macrophage | confirmed (strong) | PC, DirectLiNGAM |
| Pro-inflammatory macrophage → efferocytosis-poised macrophage | confirmed (strong) | DirectLiNGAM |
| Extracellular-matrix fibroblast → growth-factor fibroblast | confirmed (strong) | PC, DirectLiNGAM, GRaSP |
| Pro-inflammatory macrophage → alveolar-macrophage sensing | reversed (textbook GT: sensing → pro-inflammatory) | multi-method |
| Endothelial-barrier program → extracellular-matrix fibroblast | reversed (strong-evidence GT: matrix → barrier) | multi-method |
| Alveolar-macrophage sensing → CD4 activation | confirmed (supportive) | multi-method |
| Alveolar-macrophage sensing → NK cytotoxic program | confirmed (supportive) | multi-method |
| Germinal-centre B-cell program → CD8 priming | reversed | multi-method |
| Germinal-centre B-cell program → endothelial-barrier program | confirmed (supportive) | multi-method |
| Germinal-centre B-cell program → extracellular-matrix fibroblast | confirmed (supportive) | multi-method |
| CD4 activation → extracellular-matrix fibroblast | confirmed (supportive) | multi-method |
| CD4 activation → CD8 cytotoxic-T program | confirmed (textbook) | multi-method |
| CD4 activation → NK cytotoxic program | confirmed (supportive) | multi-method |
| Growth-factor fibroblast → NK cytotoxic program | novel | multi-method |
| Efferocytosis-poised macrophage → alveolar-macrophage sensing | novel | multi-method |
| Efferocytosis-poised macrophage → CD4 activation | novel | multi-method |
| Efferocytosis-poised macrophage → endothelial-barrier program | confirmed (supportive) | multi-method |
| Efferocytosis-poised macrophage → growth-factor fibroblast | confirmed (supportive) | multi-method |
| Macrophage recruitment → efferocytosis-poised macrophage | confirmed (supportive) | multi-method |
| Pro-inflammatory macrophage → germinal-centre B-cell program | novel | multi-method |
| Pro-inflammatory macrophage → CD8 priming | confirmed (supportive) | multi-method |
| Pro-inflammatory macrophage → growth-factor fibroblast | confirmed (supportive) | multi-method |
| CD8 cytotoxic-T program → NK cytotoxic program | confirmed (supportive) | multi-method |

**Table 5b.** Cohort-discriminative-program arrows in the patient-stratified JCI headline DAG (a025_bt6_bn100 union; 9 program-to-context arrows + 1 between-context artefact arrow). The 26-edge count reported in the Cross-context causal network section refers to program × program edges only and excludes both the 9 program-to-context arrows and the between-context artefact arrow below.

| Program node | Cohort context | Class (cohort the program discriminates) | One-sentence biological interpretation |
| --- | --- | --- | --- |
| Germinal-centre B-cell program | Lung cancer | Cancer-discriminative | Tertiary lymphoid structures with germinal-centre activity are a recognised feature of the lung cancer microenvironment and a candidate response-correlate for immune-checkpoint blockade. |
| CD8 priming | COVID-19 | COVID-discriminative | Antigen-presentation-driven CD8 priming is upregulated in severe COVID-19 lungs relative to normal tissue. |
| Endothelial-barrier program | COVID-19 | COVID-discriminative | Endothelial-barrier dysregulation is a hallmark of severe COVID-19 pneumonia. |
| Endothelial-barrier program | Lung cancer | Cancer-discriminative | Vascular remodelling and barrier dysregulation are also enriched in the tumour microenvironment, distinguishing it from normal tissue. |
| Efferocytosis-poised macrophage | COVID-19 | COVID-discriminative | Efferocytic clearance of apoptotic cells is elevated in lethal COVID-19 lungs as part of resolution-phase macrophage programming. |
| Pro-inflammatory macrophage | COVID-19 | COVID-discriminative | Pro-inflammatory macrophage polarisation is a defining feature of severe-COVID-19 lung pathology. |
| CD8 cytotoxic-T program | COVID-19 | COVID-discriminative | Cytotoxic CD8 effector activity is selectively elevated in COVID-19 lungs versus normal tissue. |
| NK cytotoxic program | COVID-19 | COVID-discriminative | NK-cell cytotoxic programs are enriched in severe-COVID-19 lung infiltrates. |
| NK cytotoxic program | Lung cancer | Cancer-discriminative | NK-cell cytotoxic programs are also enriched in the tumour microenvironment, partially overlapping with the COVID-19 signature. |
| Lung cancer context indicator → COVID-19 context indicator † | (between-context arrow) | n/a (not biologically interpretable) | Background knowledge forbids context-context edges for the constraint-based methods but not for DirectLiNGAM; this between-context arrow is a method artefact retained only for accounting completeness. |

† Between-context arrow is a known artefact of the one-hot JCI encoding when processed by DirectLiNGAM, which does not natively consume the edges-into-context background-knowledge constraint. It is excluded from the 26-edge program × program count in the Cross-context causal network section, from the cohort-discriminative-program total (9 program-to-context arrows), and from all downstream biological interpretation.

A direction-quality companion configuration — α = 0.025, threshold = 0.7, 200 bootstrap iterations (`a025_bt7_bn200` majority, 13 edges) — achieves 83.3% direction accuracy at evaluable recall = 0.750 with zero forbidden violations, the cleanest direction corner across the entire ablation series. Together the two configurations trace the recall–direction Pareto front of the patient-stratified pool.

GES was excluded from the patient-stratified joint-causal-inference pool because its single-run BIC-search exceeded the per-analysis wall-time budget on this cohort (the Discussion); the reported consensus on this pool is therefore based on PC, FCI, DirectLiNGAM, and GRaSP, and the majority threshold scales to ≥ 3 of 4 methods (the Multi-algorithm causal discovery section). The recall = 1.000 corner is robust to GES’s absence by construction of the union consensus: the four-method union already recovers all eight evaluable ground-truth edges (Table 5), and adding GES — which would only add edges in a union consensus, never remove them — cannot reduce evaluable recall below 1.000. Across the patient-stratified Pareto front we additionally verified that no grid point in the four-method sweep drops recall below the headline (the Pareto fronts section), and the direction-corner companion configuration `a025_bt7_bn200` majority remains a clean Pareto corner under the same four-method consensus. The recall ceiling is therefore not an artefact of GES exclusion, although the direction-accuracy and forbidden metrics at majority can in principle shift if GES were added; this is an explicit scalability caveat, addressed further in the Discussion.

The six ground-truth-matching confirmed edges (the textbook plus strong-evidence subset of Table 5) in the headline patient-stratified union DAG span four canonical immune and stromal axes: CD4 activation → CD8 priming and CD8 priming → cytotoxic effector (the textbook CD4-helper / CD8-effector cascade^38^,^39^); CD4 activation → germinal-centre B-cell formation (T-helper-driven germinal-centre formation); macrophage recruitment → pro-inflammatory macrophages and pro-inflammatory macrophages → efferocytosis-poised macrophages (sequential macrophage polarisation^43^,^44^); and extracellular-matrix fibroblasts → growth-factor fibroblasts (stromal coupling). Four additional edges carry literature-supportive orientations beyond the strict curated set (CD4 activation → CD8 cytotoxic-T program; CD4 activation → extracellular-matrix fibroblast; alveolar-macrophage sensing → NK cytotoxic program; efferocytosis-poised macrophage → growth-factor fibroblast). Of the two reversed edges, both are skeleton-correct but direction-inverted: pro-inflammatory macrophages → alveolar-macrophage sensing (textbook order is sensing → pro-inflammatory) and endothelial-barrier program → extracellular-matrix fibroblasts (strong-evidence order is matrix → barrier). No forbidden-orientation violation occurs at this configuration.

Edge classes in Table 5 are assigned against the curated directed ground-truth set defined in the Ground-truth validation section (KEGG^30^, Reactome^31^, ImmuneSigDB^32^, and textbook immunology^38–46^,^56–61^): **confirmed** = correct skeleton and direction; **reversed** = correct skeleton, inverted direction; **novel** = absent from the ground-truth set (hypothesis-generating); **forbidden** = direction contradicts a forbidden-orientation constraint.

### Cohort-discriminative-program arrows in the headline patient-stratified DAG

Beyond the 26 program × program edges, the headline patient-stratified JCI DAG contains nine arrows that connect a program node to a cohort-context indicator. Background knowledge in our implementation forbids program → context arrows for the constraint-based and score-based methods (PC, FCI, GES) but DirectLiNGAM does not natively accept this prior, so program-to-context arrows can survive into the consensus through DirectLiNGAM’s contribution. We retain them as a side-table and reinterpret them as **cohort-discriminative-program arrows**: programs whose patient-level activity is statistically informative about the patient’s cohort assignment in the joint pool. Because cohort assignment is fixed by patient identity and precedes any expression measurement, the biological reading of these arrows is necessarily that the cohort context is the upstream causal variable; the table reports them in the (program → cohort indicator) orientation in which DirectLiNGAM emits them but the operational interpretation is that the cohort context is the cohort-discriminative-program’s upstream cause.

The cohort-discriminative-program tally is congruent with prior single-cell and bulk literature: the COVID-19 cohort is selectively distinguished by activated cytotoxic CD8 / NK cells, endothelial-barrier dysregulation, pro-inflammatory and efferocytic macrophages, and CD8 priming; the lung cancer cohort is distinguished by germinal-centre B-cell activity, NK cytotoxic programs, and endothelial-barrier dysregulation. The overlap on endothelial-barrier and NK cytotoxic between the two disease cohorts is consistent with vascular and innate-cytotoxic remodelling being a shared feature of acute lung pathology and the tumour microenvironment.

The disease-extended cross-cohort JCI pool exhibits a single between-context arrow at its recall-best configuration and no surviving program ↔ context arrows in the union consensus, consistent with the disease-extended pool’s pseudobulk-averaged rows providing weaker cohort-discriminative signal than the patient-stratified pool.

The cohort-discriminative-program arrows in Table 5b directly identify which biological programs the COVID-19 stratum contributes uniquely to the patient-stratified JCI pool. Of the six ground-truth-matching confirmed edges in the headline DAG (Table 5, the textbook plus strong-evidence subset), four involve at least one program that is COVID-discriminative according to Table 5b: macrophage recruitment → pro-inflammatory macrophage, pro-inflammatory macrophage → efferocytosis-poised macrophage, CD8 priming → CD8 cytotoxic-T program, and CD4 activation → CD8 priming. The COVID stratum therefore contributes specific, non-redundant signal to two of the four canonical biological axes recovered in the headline graph (the sequential-macrophage-polarisation axis and the CD8-priming → cytotoxic-effector axis). The remaining two confirmed edges (CD4 activation → germinal-centre B-cell program; extracellular-matrix fibroblast → growth-factor fibroblast) involve programs that are not COVID-discriminative; the germinal-centre arrow is cancer-discriminative through the germinal-centre B-cell program (Table 5b) and the fibroblast arrow is broadly shared across all three cohorts. A formal leave-one-stratum-out JCI sensitivity analysis — recomputing the headline DAG with each of the three strata removed in turn — was not performed within the wall-time budget of this study and is flagged as a target for a per-stratum contribution audit in future work (the Discussion).

### Disease-extended single-cohort direction collapse — mechanism and resolution

Single-cohort causal inference on the disease-extended pooled cohorts (the COVID-extended and cancer-extended cohorts that score 26 / 34 programs on all 611 patients) recovers strong skeleton signal (Table 6) but suffers a structural direction collapse that the patient-stratified joint-causal-inference pool dissolves. Skeleton evaluable recall on the COVID-extended pooled cohort reaches 0.350 / 0.350 / 0.550 (intersection / majority / union) — a sevenfold lift over the matched COVID-19 patient-stratified cohort (0.050 majority recall, sample-size-floor-locked). The cancer-extended pooled cohort behaves similarly: 0.310 / 0.310 / 0.552 evaluable recall, a 4.5-fold lift over the lung cancer patient-stratified cohort. Direction accuracy, however, collapses: the COVID-extended pooled cohort’s majority direction accuracy is 0.143 (one of seven oriented edges correct) with a 40% forbidden-violation rate, and at union the forbidden-violation rate climbs to 60% (three of five evaluable forbidden edges violated). The cancer-extended pooled cohort performs better but is still direction-limited (majority direction accuracy = 0.333; union forbidden rate = 25%).

**Table 6.**
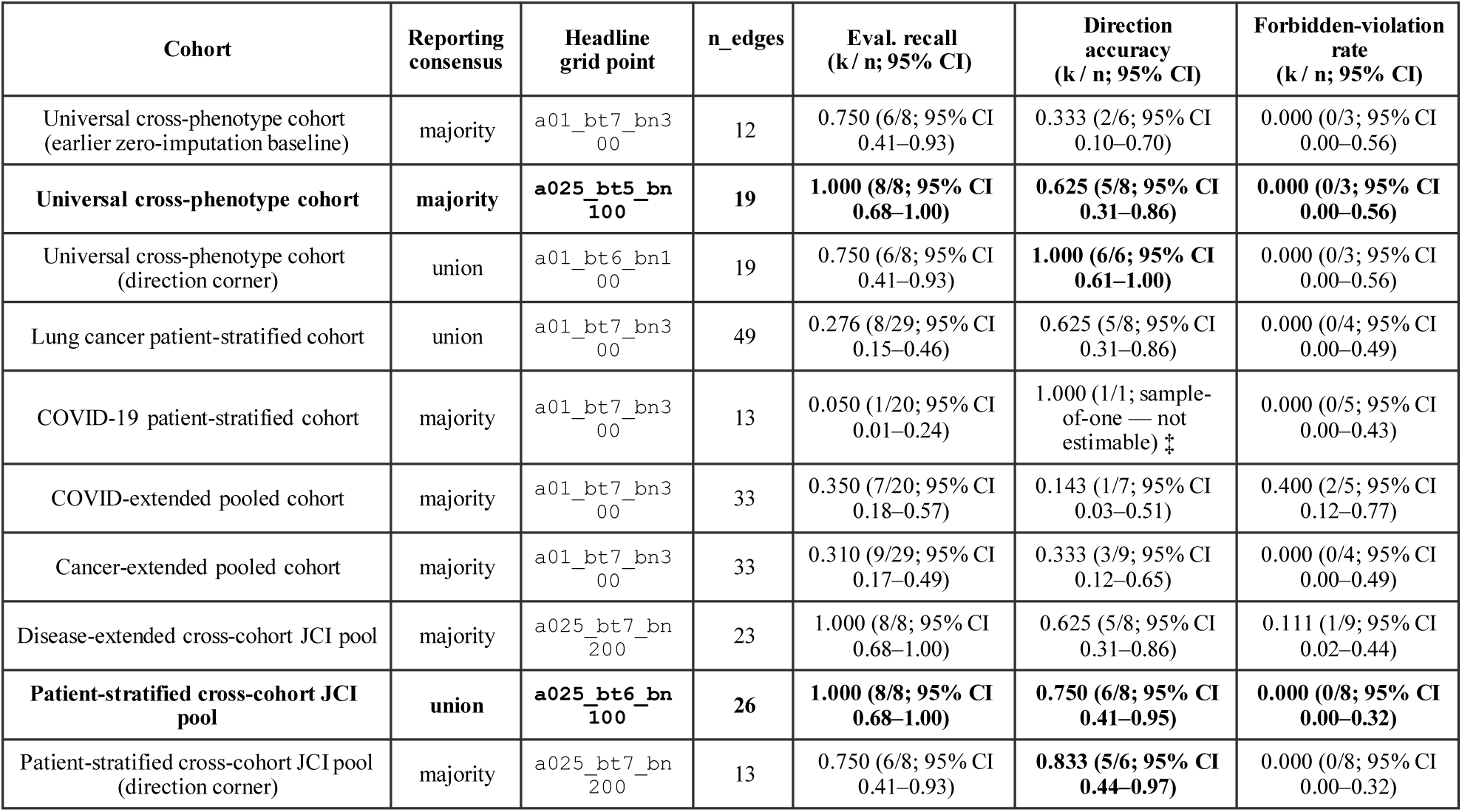
Headline validation metrics across reporting cohorts, with Wilson 95% confidence intervals on every binomial rate.

The mechanism is exposed by the contrast between the two JCI pools (Table 6, last two rows). The disease-extended cross-cohort JCI pool stacks the disease-extended pooled cohorts; despite hitting evaluable recall = 1.000 at majority and union under the systematic hyperparameter sweep with low-percentile imputation (`a025_bt7_bn200` / `a025_bt7_bn300`), it never exceeds direction accuracy = 0.625 at recall = 1.000 and carries one forbidden violation at every recall-best configuration. The patient-stratified cross-cohort JCI pool stacks strictly disjoint patient populations (no within-cohort normal-plus-disease mixing) and hits direction accuracy = 0.750 at recall = 1.000 with zero forbidden violations on the headline configuration (Figure 4). The +0.125 direction-accuracy gain and -1 forbidden-violation gain at matched recall is the quantitative evidence that the disease-extended direction collapse is caused by within-cohort pooling over heterogeneous patient populations: the cancer-extended pooled cohort mixes ∼458 non-cancer patients (370 normal + 88 COVID-19) with 153 lung cancer patients, and the COVID-extended pooled cohort mixes ∼523 non-COVID patients (370 normal + 153 cancer) with 88 COVID-19 patients. In both cases the algorithms see two superimposed latent regimes orienting average rather than disease-specific structure. The collapse is resolved at the cross-context level by joint-causal-inference pooling on patient-stratified cohorts.

**Figure 1.**
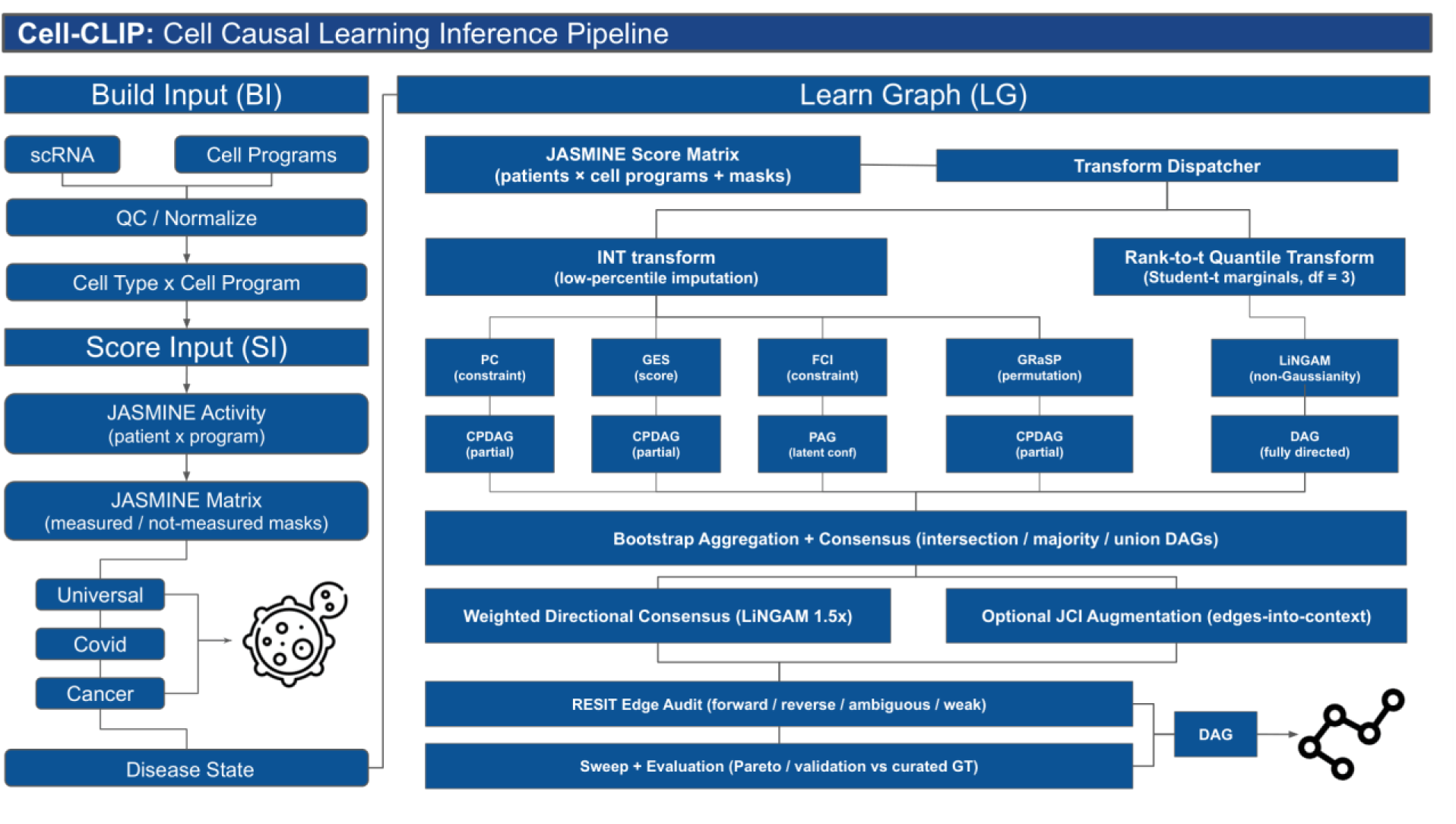
Cell-CLIP modular workflow, organised as two linked panels: a **Build-Input** panel (left) and a **Learn-Graph** panel (right). The Build-Input panel ingests per-phenotype scRNA-seq from the three contributing cohorts (universal cross-phenotype, COVID-19, lung cancer) and, after quality-control and library-size normalisation, scores cell programs with cell-type-aware mask-aware JASMINE — producing a per-patient × per-cell-type-program activity matrix together with an explicit missing-value mask that distinguishes “not measured” (off-context or low cell count) from “low activity”. The Learn-Graph panel applies a per-algorithm transform dispatcher (low-percentile imputation in raw activity space followed by the inverse-normal transform for PC, GES, FCI, GRaSP; the same low-imputed matrix followed by a rank-to-Student-t (df = 3) transform for DirectLiNGAM), runs the five causal-discovery algorithms in parallel under bootstrap resampling, aggregates per-algorithm directed edges via the two-phase weighted consensus to produce intersection, majority, and union CPDAG/PAG/DAG outputs, optionally augments via Joint Causal Inference with edges-into-context background knowledge, and finally annotates each consensus edge with the RESIT residual-independence audit (forward / reverse / ambiguous / weak) before evaluation against curated ground truth on a systematic three-by-three-by-three hyperparameter grid. The patient icon between the per-phenotype source boxes and the disease-state row denotes the **patient-cohort** layer, i.e., the patient-level partition that subsequent JCI pooling treats as the context variable.

**Figure 2.**
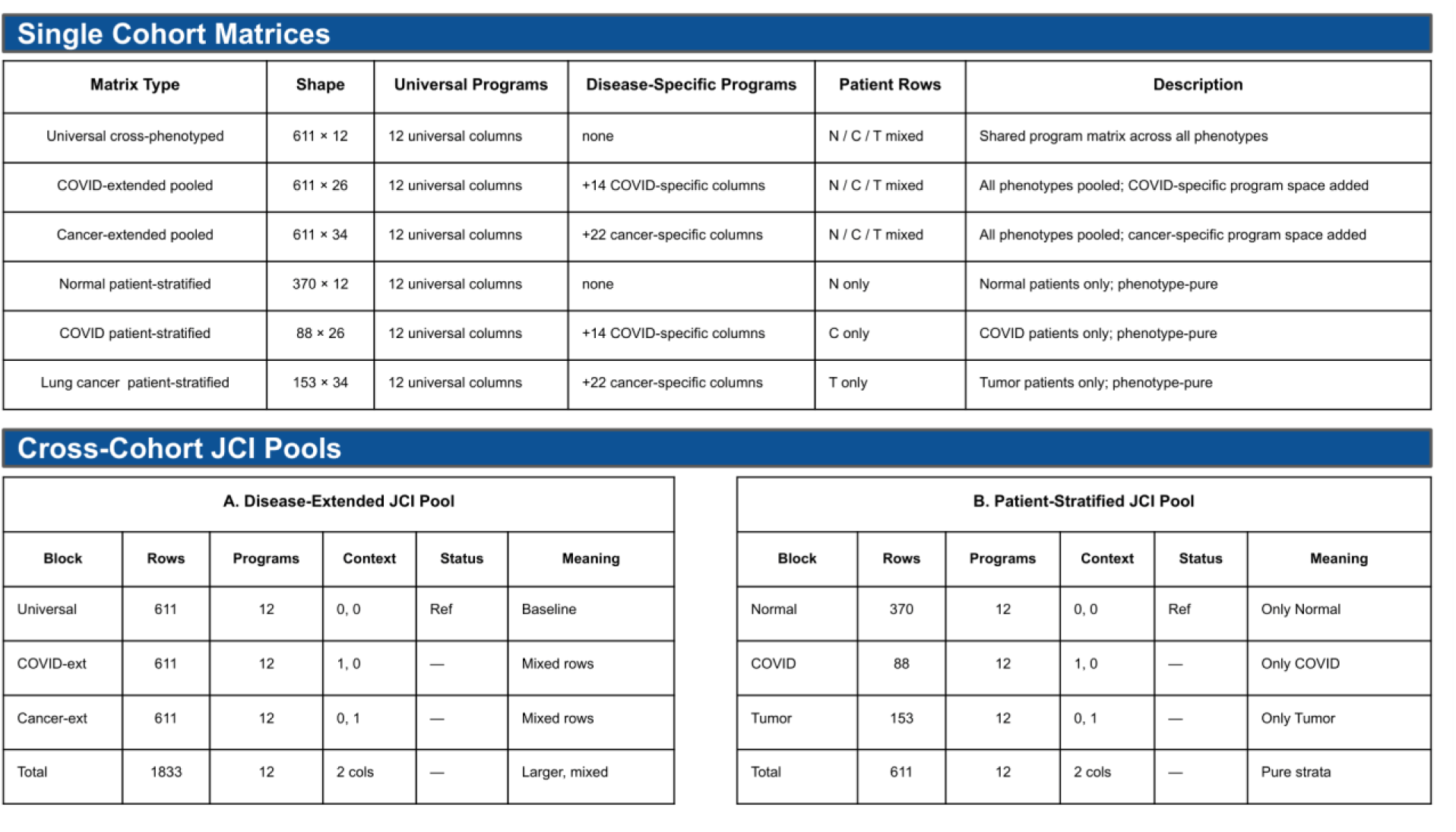
Single-cohort matrices and joint-causal-inference (JCI) pools. The top table summarises the five single-cohort matrices and the normal-tissue reference cohort (cohort label, programs, patients, mask-aware fill rate, disease state). The bottom table summarises the two JCI pools, with columns for the constituent cohorts, the dropped reference indicator, and the total augmented row count. The patient-stratified JCI pool is highlighted as the headline cross-context graph. *Footnote — JCI pools use the 12 universal program columns only; the disease-specific COVID and cancer program columns shown in Table 1 are not carried into either pool, because cross-context inference requires a shared program set across all contributing cohorts*.

**Figure 3.**
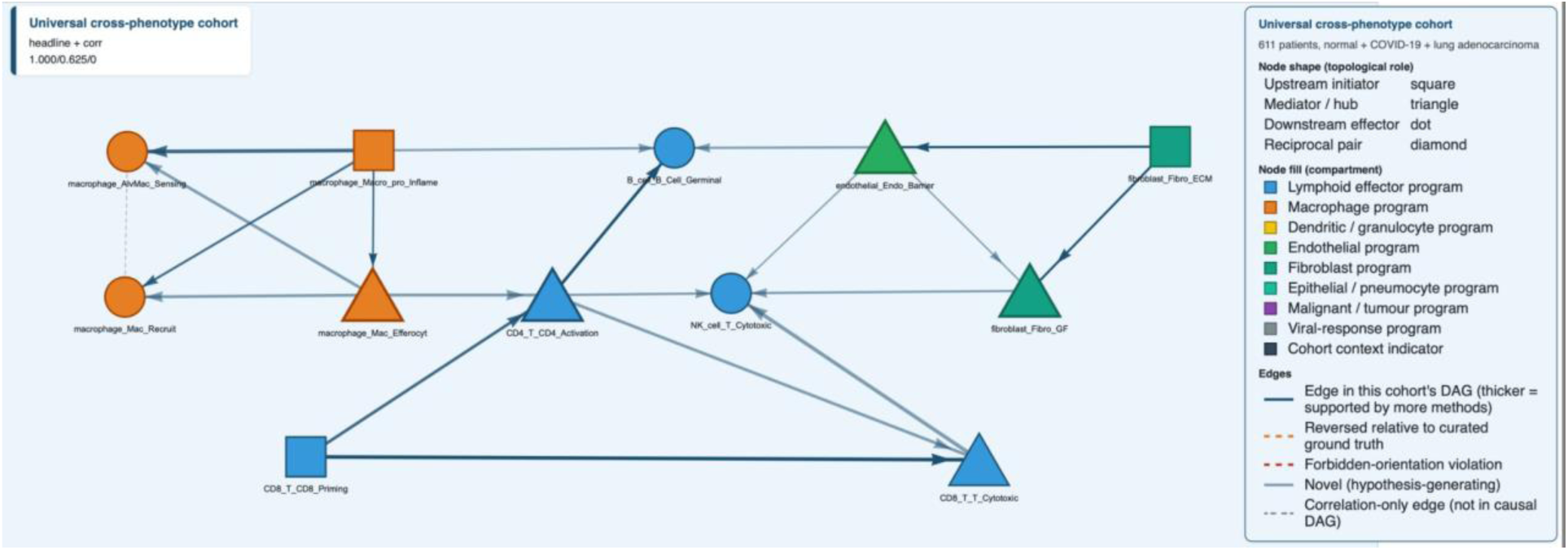
Universal cross-phenotype headline majority directed acyclic graph (low-percentile imputation, hyperparameter setting α = 0.025, bootstrap threshold = 0.5, 100 bootstrap iterations; 19 edges). Node shape encodes the program’s topological role inside the graph (square = upstream initiator with no incoming edges, triangle = mediator hub with both incoming and outgoing edges, circle = downstream effector with no outgoing edges); cross-compartment hubs are drawn with thicker borders. Node fill encodes the cell-type compartment (lymphoid effector in blue, macrophage in orange, dendritic / granulocyte in yellow, endothelial in green, fibroblast in teal-green, epithelial in cyan, malignant in purple, viral-response in grey). Edge colour encodes edge class: deep navy at full opacity for **confirmed** edges (direction matches curated ground truth), deep navy at 55% opacity for **novel** edges (not in ground truth, hypothesis-generating), dashed orange for **reversed** edges (correct skeleton, inverted direction relative to ground truth), dashed red for **forbidden** edges (direction contradicts a forbidden-orientation constraint). Edge thickness encodes the number of contributing causal-discovery algorithms (range 1 to 5).

**Figure 4.**
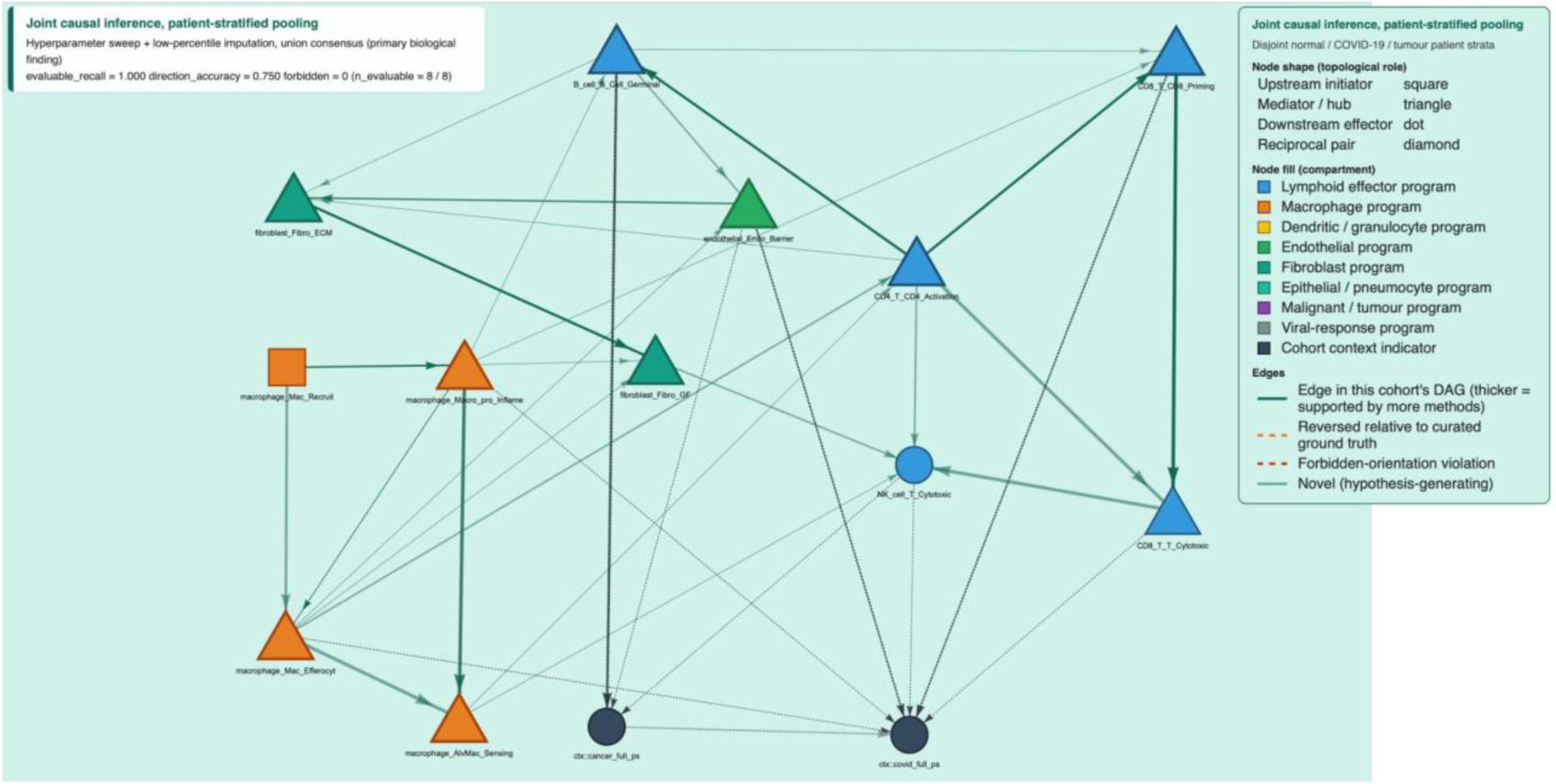
Patient-stratified cross-context joint-causal-inference union directed acyclic graph (hyperparameter setting α = 0.025, bootstrap threshold = 0.6, 100 bootstrap iterations; 26 program-by-program edges plus 9 program-to-context-indicator arrows and 1 context-to-context arrow; evaluable recall = 1.000, direction accuracy = 0.750, zero forbidden-orientation violations). Node shape and fill encode topological role and compartment as in Figure 3; the two cohort-context indicators (COVID-19 patient stratum, lung-cancer patient stratum; the normal-tissue stratum is the dropped reference) are rendered in dark slate as dedicated context nodes. Edge colour is the patient-stratified joint-causal-inference accent (dark teal-green) for confirmed and novel orientations, dashed orange for reversed orientations, dashed red for forbidden-orientation violations. Cohort-discriminative-program arrows from the Cohort-discriminative-program arrows section and Table 5b are rendered as fine dashed slate lines pointing from biological programs into the context indicators.

**Figure 5.**
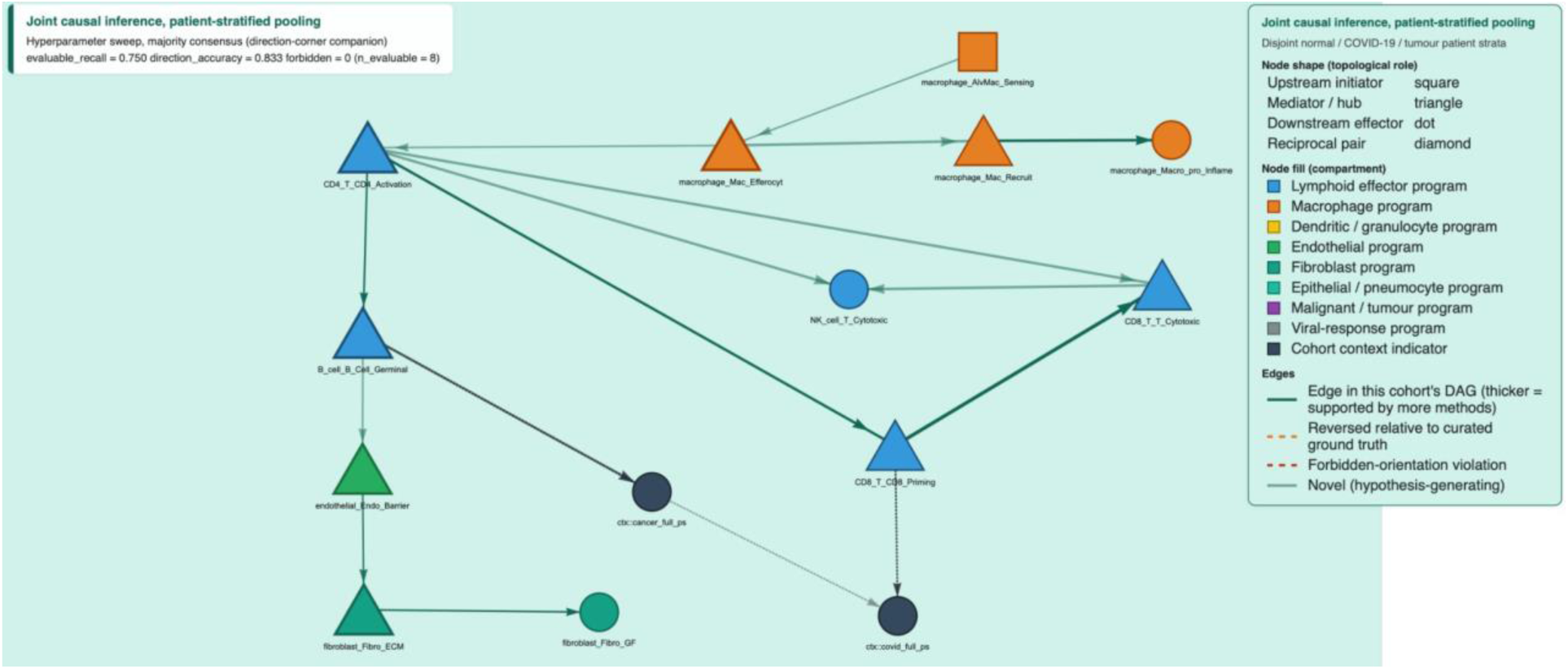
Patient-stratified cross-context joint-causal-inference direction-corner majority directed acyclic graph (hyperparameter setting α = 0.025, bootstrap threshold = 0.7, 200 bootstrap iterations; 13 edges; evaluable recall = 0.750, direction accuracy = 0.833, zero forbidden-orientation violations). Same shape, fill, and edge encoding as Figure 4. This direction-corner companion graph is sparser than Figure 4 but achieves the highest direction accuracy across the entire ablation series at non-trivial recall.

**Figure 6.**
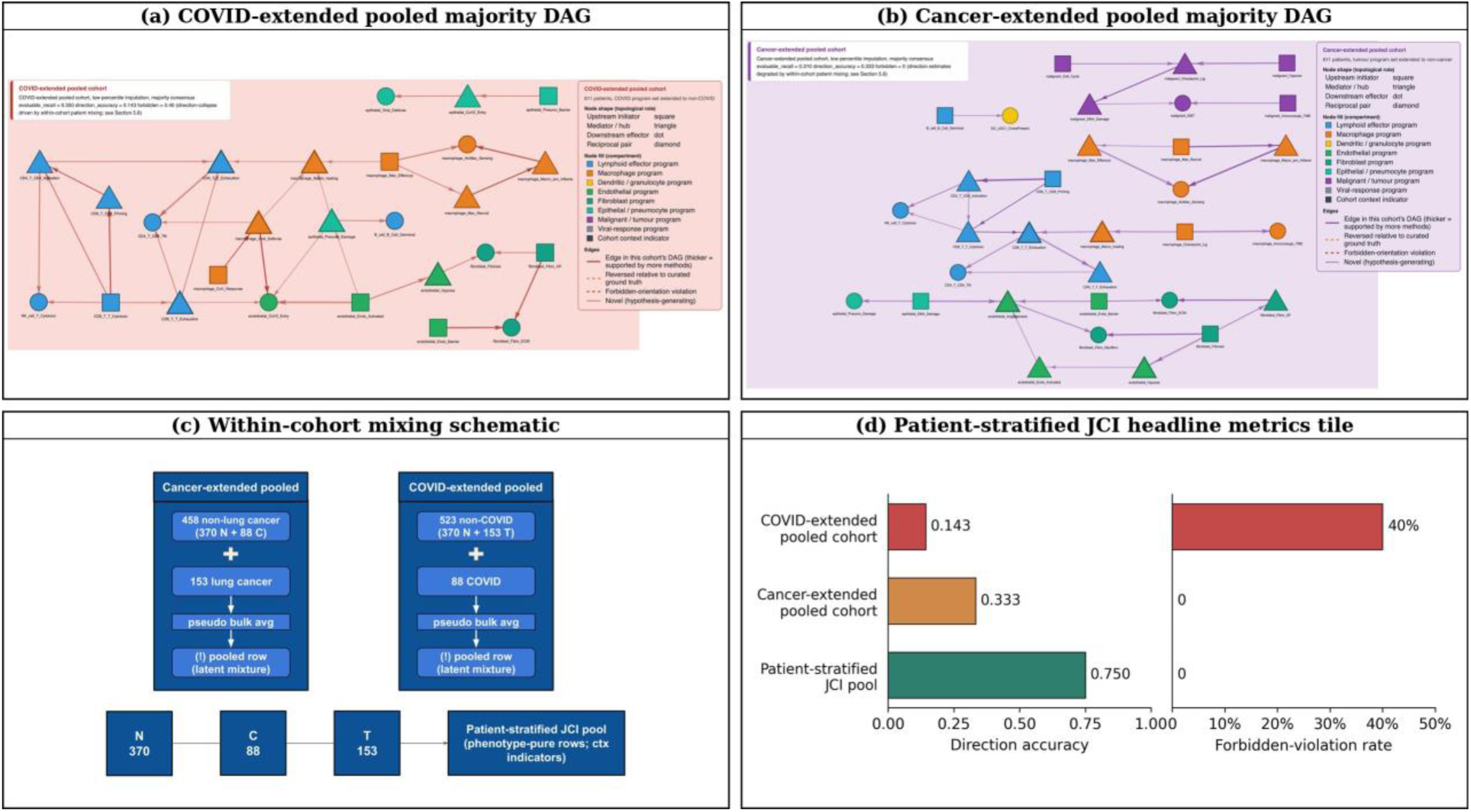
Direction collapse on the disease-extended single-cohort matrices and its resolution by patient-stratified joint-causal-inference pooling. **(a)** COVID-extended pooled cohort majority directed acyclic graph (33 edges; evaluable recall = 0.350, direction accuracy = 0.143, forbidden-violation rate = 40%) with two highlighted forbidden orientations (coronavirus-host-response program oriented as upstream of the viral-defence program; viral-defence program oriented as upstream of the SARS-CoV-2 cell-entry program). **(b)** Cancer-extended pooled cohort majority directed acyclic graph (33 edges; evaluable recall = 0.310, direction accuracy = 0.333, forbidden-violation rate = 0). **(c)** Schematic of the within-cohort mixing mechanism. Each disease-extended pooled cohort averages a large non-disease majority of patients (approximately 458 non-cancer patients in the cancer-extended pool — 370 normal and 88 COVID-19; approximately 523 non-COVID patients in the COVID-extended pool — 370 normal and 153 lung cancer) together with the much smaller disease subset, so that the joint causal-discovery problem implicitly superimposes two patient sub-populations with potentially distinct causal regimes; the panel illustrates this superposition and shows that pre-stratifying patients by disease state before pooling — the construction used by the patient-stratified joint-causal-inference pool (Figure 4) — avoids it. **(d)** Quantitative summary tile contrasting the disease-extended cross-cohort joint-causal-inference pool with the patient-stratified cross-cohort joint-causal-inference pool at their respective headline configurations on the three primary validation metrics (evaluable recall, direction accuracy, and forbidden-violation rate), showing the +0.125 direction-accuracy gain and -1 forbidden-violation gain at matched evaluable recall = 1.000 that constitutes the quantitative evidence for the mixing-induced direction collapse described above and in Table 6.

**Figure 7.**
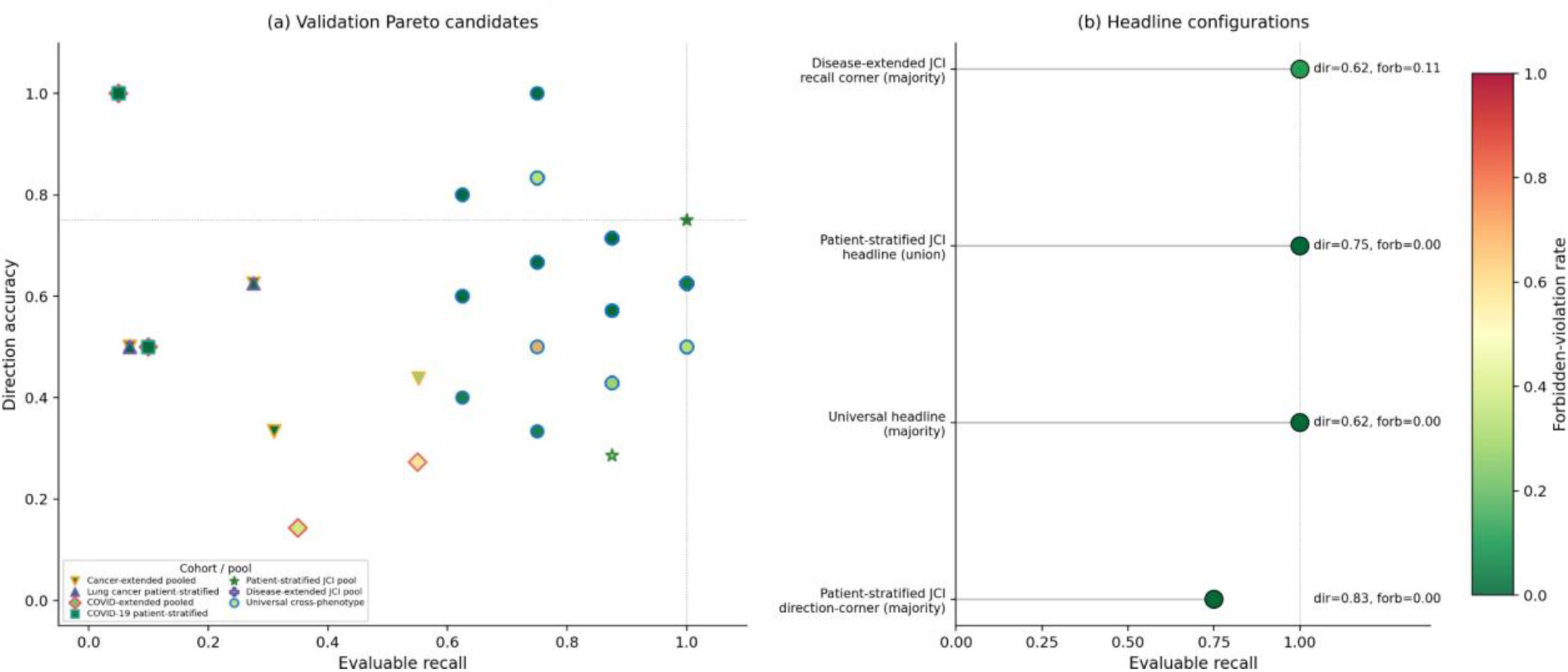
Validation Pareto fronts across cohorts shown as two linked panels produced from the systematic hyperparameter grid: scatter of all cohort × consensus × grid-point validations (markers encode cohort; marker fill colour maps to forbidden-orientation violation rate); and a condensed headline-configuration summary listing direction accuracy and forbidden-violation rates for the four labelled flagship configurations. The COVID-extended pooled cohort concentrates in the high-evaluable-recall / low-direction-accuracy quadrant of panel (a), mirroring the pooled-cohort direction collapse described in the Disease-extended single-cohort direction collapse section.

### Patient-stratified single-cohort companions

Cell-CLIP also produces single-cohort patient-stratified graphs as companions to the cross-context headline. The lung cancer patient-stratified cohort under low-percentile imputation (`a01_bt7_bn300` union, 49 edges; Table 6) recovers 0.276 evaluable recall — matching the recall ceiling obtained by the earlier zero-imputation baseline configuration on this cohort (Table 8) and constituting the only single-cohort tumour evaluable-recall ceiling reached by any pipeline configuration in the ablation series — at direction accuracy = 0.625, the strongest single-cohort tumour direction accuracy seen in the ablation series. Majority consensus on the same cohort regresses (evaluable recall 0.069), so we report the lung cancer patient-stratified cohort at union only. The COVID-19 patient-stratified cohort floor-locks at 0.050 evaluable recall under every imputation strategy — its 88-patient × 26-program ratio (n / p ≈ 3.4) is below Fisher’s-Z power for the relevant edge densities — but the COVID-relevant edges are recovered when this cohort is contributed as a context to the patient-stratified JCI pool (the Cross-context causal network section).

### Validation against curated ground truth across all reporting cohorts

Table 6 reports the headline evaluable recall, direction accuracy, and forbidden-violation rate for every reporting cohort, alongside a comparable earlier-baseline row (the zero-imputation baseline configuration of the pipeline, defined in the Sensitivity analysis section) where available. Three configurations achieve evaluable recall = 1.000: the universal cross-phenotype cohort under the systematic hyperparameter sweep with low-percentile imputation (majority and union), the disease-extended cross-cohort JCI pool under the same procedure (majority and union, multiple grid points), and the patient-stratified cross-cohort JCI pool (union headline). Of these, only the patient-stratified JCI pool simultaneously achieves direction accuracy ≥ 0.750 and zero forbidden violations.

Bold rows are the primary single-cohort and cross-context biological findings. All intervals are Wilson 95% confidence intervals (Methods §“Ground-truth validation”); given small evaluable ground-truth denominators (8 ≤ *n* ≤ 29), point-estimate gaps between rows are typically smaller than their respective intervals and should be interpreted as direction-of-effect rather than statistically significant differences. ‡ The COVID-19 patient-stratified cohort recovers only one oriented evaluable edge at this configuration; a sample-of-one direction accuracy is reported here for completeness but is not interpretable as a reliable estimate, and we therefore report this cohort primarily as a context contributor to the patient-stratified joint-causal-inference pool rather than as a stand-alone cohort.

### Pareto fronts across the hyperparameter sweep

For each headline cohort, the full hyperparameter sweep produces a recall × direction × forbidden Pareto front (Table 7). On the universal cross-phenotype cohort, three grid points achieve evaluable recall = 1.000 with zero forbidden violations (`a025_bt5_bn100` majority, `a05_bt5_bn100` intersection, `a05_bt7_bn200` union; all at direction accuracy = 0.625), and two grid points achieve direction accuracy = 1.000 at evaluable recall = 0.750 with zero forbidden violations (`a01_bt6_bn100` union, `a025_bt6_bn300` union). On the patient-stratified JCI pool, 23 of 27 majority grid points carry zero forbidden violations and all 27 intersection grid points carry zero forbidden violations — the cleanest forbidden record of any sweep we have validated. On the disease-extended JCI pool, the recall-maximising regime hits evaluable recall = 1.000 across most majority grid points but always carries at least one forbidden violation, consistent with the disease-extended mixing mechanism (the Disease-extended direction collapse section). Reporting per-cohort Pareto fronts rather than a single best configuration exposes the recall–direction trade-off explicitly and justifies the cohort-specific choice of consensus rule (intersection or majority for the universal cross-phenotype cohort; union for the lung cancer patient-stratified cohort; union for the patient-stratified JCI pool’s headline; majority for both JCI pools in their direction-corner roles).

**Table 7.** Pareto fronts under the systematic hyperparameter sweep with low-percentile imputation.

| Cohort | Recall corner (recall $\geq 0.875$) | Direction corner (dir $\geq 0.833$) | Notes |
| --- | --- | --- | --- |
| Universal cross-phenotype cohort | 3 grid points @ recall = 1.000 / dir = 0.625 / forbidden = 0 | 2 grid points @ recall = 0.750 / dir = 1.000 / forbidden = 0 | clean Pareto, both corners reachable |
| Disease-extended cross-cohort JCI pool | most of 27 majority grid points @ recall = 1.000 / dir = 0.625–0.833 / forbidden = 1 | 0 grid points reach (dir $\geq 0.833$ + forbidden = 0); best direction at forbidden = 0 is 0.625 | mixing-induced forbidden floor |
| Patient-stratified cross-cohort JCI pool | 1 grid point @ recall = 1.000 / dir = 0.750 / forbidden = 0 (headline); 4 grid points @ recall = 0.875 / dir = 0.714 / forbidden = 0 | 1 grid point @ recall = 0.750 / dir = 0.833 / forbidden = 0 | cleanest grid; both corners reachable |

### Sensitivity analysis: imputation, DirectLiNGAM weighting, and consensus rule

We performed an ablation series across five single cohorts and five methodological variants to isolate which methodological levers are responsible for the headline numbers (Table 8). The five variants are defined as follows and are referred to consistently by these descriptive names throughout the rest of the paper:

- **Zero-imputation baseline** — the earlier baseline configuration of the pipeline: missing entries imputed as zero in the raw JASMINE activity space, with no mask-aware handling and no per-algorithm transform dispatcher (the four linear-Gaussian methods and DirectLiNGAM all receive the same column-wise inverse-normal-transformed matrix).
- **Median-imputation baseline** — mask-aware JASMINE scoring (the Mask-aware JASMINE scoring section) combined with median imputation of missing entries on the inverse-normal-transform (INT) scale; the per-algorithm transform dispatcher is in place but low-percentile imputation is not yet used.
- **DirectLiNGAM-direction-downgrade hybrid** — the median-imputation baseline with two further modifications combined into a single sensitivity variant: (i) the DirectLiNGAM direction weight is reduced from 1.5× back to 1.0× (the Two-phase weighted directional consensus section), and (ii) a column-wise fallback rule replaces the rank-to-Student-*t* DirectLiNGAM input with the inverse-normal-transformed input on columns whose mask-aware fill rate falls below a pre-set low-fill threshold.
- **Single-configuration low-percentile imputation** — low-percentile imputation in raw activity space (the Low-percentile imputation section) run at a single representative hyperparameter setting, without a full grid sweep.
- **Systematic hyperparameter sweep with low-percentile imputation** — low-percentile imputation combined with the full three-by-three-by-three hyperparameter grid (the Hyperparameter sweep section); the production pipeline reported throughout this paper.

**Table 8.**
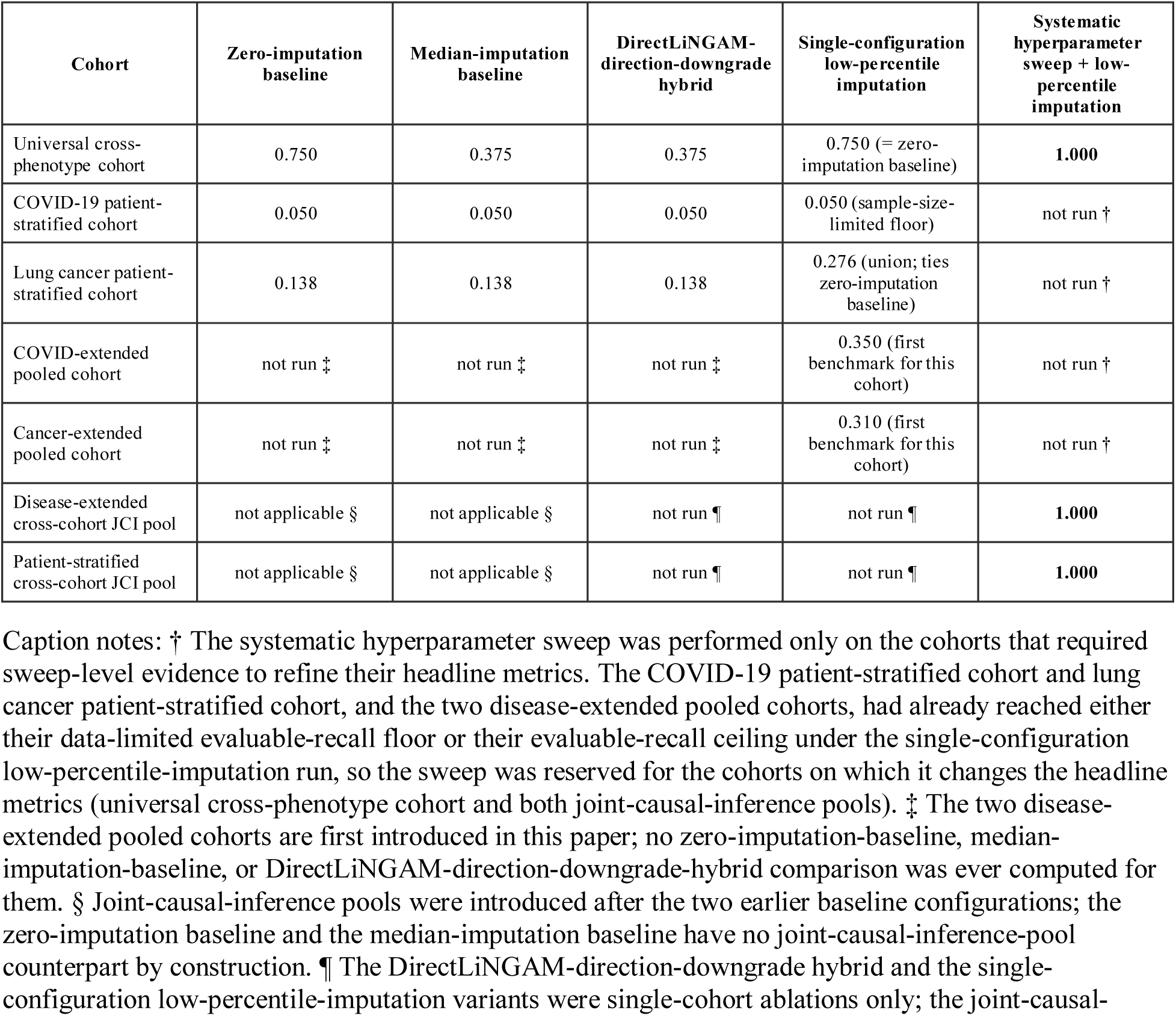
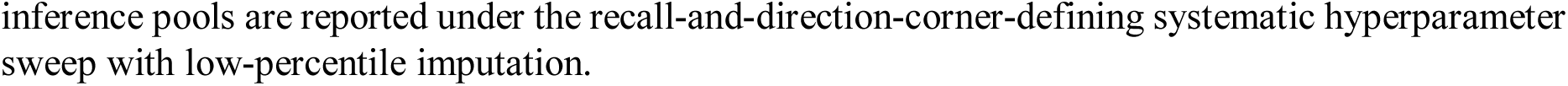
Ablation matrix: evaluable recall at the recall-best configuration. Variants are defined in the Sensitivity analysis section. Cells marked *not run* were not executed for that combination of cohort × variant; reasons are given in the table caption.

| Cohort | Zero-imputation baseline | Median-imputation baseline | DirectLiNGAM-direction-downgrade hybrid | Single-configuration low-percentile imputation | Systematic hyperparameter sweep + low-percentile imputation |
| --- | --- | --- | --- | --- | --- |
| Universal cross-phenotype cohort | 0.750 | 0.375 | 0.375 | 0.750 (= zero-imputation baseline) | **1.000** |
| COVID-19 patient-stratified cohort | 0.050 | 0.050 | 0.050 | 0.050 (sample-size-limited floor) | not run † |
| Lung cancer patient-stratified cohort | 0.138 | 0.138 | 0.138 | 0.276 (union; ties zero-imputation baseline) | not run † |
| COVID-extended pooled cohort | not run ‡ | not run ‡ | not run ‡ | 0.350 (first benchmark for this cohort) | not run † |
| Cancer-extended pooled cohort | not run ‡ | not run ‡ | not run ‡ | 0.310 (first benchmark for this cohort) | not run † |
| Disease-extended cross-cohort JCI pool | not applicable § | not applicable § | not run ¶ | not run ¶ | **1.000** |
| Patient-stratified cross-cohort JCI pool | not applicable § | not applicable § | not run ¶ | not run ¶ | **1.000** |

Caption notes: † The systematic hyperparameter sweep was performed only on the cohorts that required sweep-level evidence to refine their headline metrics. The COVID-19 patient-stratified cohort and lung cancer patient-stratified cohort, and the two disease-extended pooled cohorts, had already reached either their data-limited evaluable-recall floor or their evaluable-recall ceiling under the single-configuration low-percentile-imputation run, so the sweep was reserved for the cohorts on which it changes the headline metrics (universal cross-phenotype cohort and both joint-causal-inference pools). ‡ The two disease-extended pooled cohorts are first introduced in this paper; no zero-imputation-baseline, median-imputation-baseline, or DirectLiNGAM-direction-downgrade-hybrid comparison was ever computed for them. § Joint-causal-inference pools were introduced after the two earlier baseline configurations; the zero-imputation baseline and the median-imputation baseline have no joint-causal-inference-pool counterpart by construction. ¶ The DirectLiNGAM-direction-downgrade hybrid and the single-configuration low-percentile-imputation variants were single-cohort ablations only; the joint-causal-
inference pools are reported under the recall-and-direction-corner-defining systematic hyperparameter sweep with low-percentile imputation.

The key findings are: **(1) low-percentile imputation is the dominant lever for universal cross-phenotype recall.** Moving from the median-imputation baseline to the DirectLiNGAM-direction-downgrade hybrid yields no change in evaluable recall on the universal cross-phenotype cohort (Δ = 0.000), whereas moving from the median-imputation baseline to single-configuration low-percentile imputation recovers the evaluable recall of the earlier zero-imputation baseline (0.375 → 0.750), and the systematic hyperparameter sweep with low-percentile imputation exceeds it (0.750 → 1.000). **(2) The DirectLiNGAM-direction-downgrade hybrid failed wherever it was run.** It reduced the lung cancer patient-stratified majority direction accuracy from 0.500 to 0.250 and produced no improvement on the other four single cohorts; we therefore drop it from the production pipeline. Because this variant bundles a direction-weight downgrade with an inverse-normal-transform fallback rule, it cannot disentangle whether the regression is driven by the direction weight, the fallback rule, or their interaction; an isolated direction-weight comparison is flagged for future sensitivity work (the Discussion). **(3) The systematic hyperparameter sweep with low-percentile imputation runs faster than the equivalent sweep on the median-imputation baseline** because the low-percentile imputation step makes both the Fisher’s-Z conditional-independence tests and the per-bootstrap re-estimations converge faster (universal grid −9%, disease-extended JCI grid −40%). **(4) Cohort × consensus interactions are real**: the universal cross-phenotype cohort is best at majority; the lung cancer patient-stratified cohort is best at union; the COVID-19 patient-stratified cohort sits at the sample-size-limited evaluable-recall floor regardless of consensus rule and is reported as a context contributor to the patient-stratified joint-causal-inference pool rather than as a stand-alone graph; the disease-extended JCI pool is reported at majority for the recall-best configurations; the patient-stratified JCI pool’s headline is at union; and its high-direction-accuracy companion is at majority.

## Discussion

We present Cell-CLIP, a multi-algorithm, mask-aware, joint-causal-inference-pooled causal-discovery framework that infers directed cell-program interaction networks from patient-level scRNA-seq atlases. The key methodological contributions are: (i) a mask-aware JASMINE scoring layer that distinguishes “not measured” from “low activity”; (ii) a low-percentile imputation policy in raw activity space that preserves the absent-cell-type signal across the rank transform; (iii) a per-algorithm transform dispatcher that gives each method an input matched to its identifiability assumption; (iv) a weighted two-phase consensus that promotes DirectLiNGAM’s direction vote 1.5× to reflect its unique ability to break Markov-equivalence-class ties; (v) a joint-causal-inference pooling layer that combines patient cohorts via context indicators with edges-into-context background knowledge; (vi) a RESIT residual-independence direction audit that annotates each consensus edge with forward / reverse / ambiguous / weak; and (vii) a systematic three-by-three-by-three hyperparameter sweep with explicit per-cohort Pareto-front reporting.

Our primary biological finding is the patient-stratified cross-cohort joint-causal-inference graph, which simultaneously recovers all eight evaluable curated ground-truth edges, achieves 75% direction accuracy on those edges, and produces zero forbidden-orientation violations against eight evaluable forbidden orientations — the only configuration in our ablation series spanning seven cohort/pool configurations, 27 hyperparameter settings, and three consensus types to reach all three optimal Pareto corners. The single-cohort universal cross-phenotype graph independently reaches the recall = 1.000 corner under the same hyperparameter sweep with low-percentile imputation, with three grid points achieving direction accuracy = 0.625 and zero forbidden violations and two additional grid points achieving direction accuracy = 1.000 at recall = 0.750 with zero forbidden violations. Both findings exceed the earlier zero-imputation baseline configuration of the pipeline (defined in the Sensitivity analysis section) on every individual metric.

A second, methodologically important contribution is the **disease-extended direction-collapse finding**. Single-cohort causal inference on the disease-extended pooled cohorts recovers 5–7× more skeleton edges than the matched patient-stratified cohorts but suffers a structural direction collapse — the COVID-extended pooled cohort’s majority direction accuracy is 0.143 with a 40% forbidden-violation rate, and at union the forbidden rate climbs to 60%. We show that this collapse is not a methodological failure of the pipeline but a structural consequence of within-cohort pooling over heterogeneous normal-plus-disease patient populations: the cancer-extended pooled cohort mixes ∼458 non-cancer patients (370 normal + 88 COVID-19) with 153 lung cancer patients, while the COVID-extended pooled cohort mixes ∼523 non-COVID patients (370 normal + 153 cancer) with 88 COVID-19 patients. In both cases the algorithms see two superimposed latent causal regimes, and joint-causal-inference pooling on patient-stratified cohorts dissolves the collapse without modifying any algorithm. The +0.125 direction-accuracy gain and -1 forbidden-violation gain between the disease-extended pool and the patient-stratified pool at matched evaluable recall = 1.000 is the quantitative evidence for this mechanism. This finding generalises beyond this dataset: any multi-cohort study where each cohort itself averages over heterogeneous patient subpopulations should expect a within-cohort mixing-induced direction collapse, and the canonical resolution is to pre-stratify patients before pooling.

A third contribution is **methodological honesty about ground-truth coverage and sample size**. The COVID-19 patient-stratified cohort (88 patients × 26 programs, n / p ≈ 3.4) sits at the evaluable-recall floor under every imputation strategy, and we report this directly rather than mask it: 88-patient cohorts are below Fisher’s-Z power for the relevant partial-correlation densities, and no imputation policy or hyperparameter choice changes that. The COVID-relevant edges are nonetheless recovered at the cross-context level when this cohort is contributed to the patient-stratified joint-causal-inference pool — a positive finding that joint-causal-inference pooling rescues otherwise-power-limited cohorts.

Several methodological points deserve emphasis. First, the per-algorithm transform dispatcher is a small structural change with a measurable effect: feeding DirectLiNGAM the same standard-normal INT as the linear-Gaussian methods erases the non-Gaussianity its pairwise independence test requires, and the rank-to-Student-t (df = 3) transform restores the heavy-tailed marginals (median kurtosis ≈ 3.8 versus 0 for INT) that LiNGAM exploits for direction identifiability. Second, the weighted directional consensus reflects an asymmetry of identifiability: PC, GES, FCI, and GRaSP cannot orient edges within a Markov-equivalence class, so their direction votes within those classes are arbitrary tie-breakers rather than evidence; DirectLiNGAM is the only method that can identify direction within such classes under the linear-non-Gaussian assumption. The 1.5× direction-weight promotion is a conservative reflection of that asymmetry, chosen empirically rather than from a parameter scan, and is used uniformly across all reported runs (the Two-phase weighted directional consensus section); the DirectLiNGAM-direction-downgrade hybrid sensitivity variant (the Sensitivity analysis section), which combined a direction-weight downgrade from 1.5× back to 1.0× with an inverse-normal-transform fallback rule applied to low-fill columns, produced no improvement and is dropped from the production pipeline. Third, the RESIT audit is a confidence statement, not a correction. Reporting forward / reverse / ambiguous / weak counts per cohort gives reviewers direct visibility into how much of the directed graph is independently supported by an additive-noise residual-independence test versus imposed by algorithm conventions; a substantial fraction of edges in the noisier cohorts are direction-unidentifiable from residuals (the Universal causal network section), and only a smaller core is cross-validated by independence tests.

End-to-end execution across the four primary cohorts and two joint-causal-inference pools required approximately 41.5 h of single-GPU/CPU compute (Google Colab A100 instance), with the patient-stratified joint-causal-inference pool’s full hyperparameter sweep completing in 3.4 h. A small number of analyses exceeded the 10-hour single-analysis wall-time budget we used as a per-cohort ceiling — the cancer-extended pooled cohort, the lung cancer patient-stratified cohort under cell-type-agnostic aggregation, and the patient-stratified joint-causal-inference pool’s full hyperparameter sweep — and are discussed alongside the resulting GES-exclusion consequences in the Limitations subsection below.

Future work falls into five categories. First, experimental validation of the highest-confidence predictions, particularly the novel efferocytosis-driven CD4-activation edge together with its literature-supported partner efferocytosis-driven macrophage-recruitment edge (both shared between the universal and patient-stratified joint-causal-inference graphs and jointly constituting a candidate efferocytosis-driven myeloid-lymphoid feedback signal), and the cohort-discriminative-program arrows in Table 5b that constitute the framework’s most direct disease-specific predictions. Second, expanding the framework to additional tissues and disease states with patient-stratified cohorts so that the joint-causal-inference pooling layer can be exercised across more heterogeneous biology. Third, a targeted LiNGAM-direction audit on the disease-extended single-cohort forbidden edges to characterise whether a small set of LiNGAM-driven flips drives the within-cohort mixing collapse — if so, a per-cohort LiNGAM-weight schedule may recover usable disease-extended single-cohort direction accuracy without requiring patient-stratification. Fourth, an isolated DirectLiNGAM direction-weight sensitivity scan that holds the rest of the production pipeline fixed and varies only the DirectLiNGAM direction weight (1.0×, 1.5×, 2.0×) on the universal cross-phenotype cohort and the patient-stratified joint-causal-inference pool, to disentangle the 1.5× direction-weight promotion from the inverse-normal-transform fallback component of the DirectLiNGAM-direction-downgrade hybrid (the Two-phase weighted directional consensus section, the Sensitivity analysis section). Fifth, a leave-one-stratum-out joint-causal-inference sensitivity analysis that recomputes the headline patient-stratified DAG with each of the three strata (normal, COVID-19, lung cancer) removed in turn, to quantify each stratum’s unique contribution to the eight evaluable confirmed-edge set and to the cohort-discriminative-program arrows in Table 5b.

## Limitations

- **Ground-truth denominator is small.** Across the seven reporting cohorts, 8 ≤ *n*_eval ≤ 29 curated ground-truth edges contribute to the evaluable-recall and direction-accuracy metrics. Wilson 95% confidence intervals (Methods §“Ground-truth validation”; Table 6) are correspondingly wide, and the point-estimate improvements of the production pipeline over the earlier zero-imputation baseline lie within those intervals; the headline numbers should therefore be read as direction-of-effect rather than as confirmatory significance at conventional levels.
- **No empirical correlation baseline on this dataset.** The Introduction’s claim that correlation- and co-expression-based methods cannot disentangle direct from confounded or mediated edges is grounded in the established causal-inference literature^15–18^,^50^. An explicit WGCNA-on-program-scores and GSVA-with-partial-correlation comparison on the same mask-aware JASMINE input matrix would convert this literature claim into a result on this dataset; that comparison was scoped for the present manuscript, deferred for tractability, and is planned for a dedicated follow-up methodological note rather than reported here.
- **DirectLiNGAM 1.5× direction-weight is an unscanned engineering choice.** The 1.5× value reflects the structural asymmetry that DirectLiNGAM is the only ensemble method that can break Markov-equivalence-class ties under the linear-non-Gaussian assumption; it was not selected by an isolated {1.0×, 1.5×, 2.0×} parameter scan. The DirectLiNGAM-direction-downgrade hybrid sensitivity variant (the Sensitivity analysis section) bundles a direction-weight downgrade with an inverse-normal-transform fallback rule and therefore does not isolate the weight choice; an isolated scan is listed in the Future-work paragraph above.
- **GES exclusion on the densest cohorts is a compute-budget consequence.** Consensus on the lung-cancer patient-stratified cohort, the cancer-extended pooled cohort, and the patient-stratified joint-causal-inference pool is a four-method consensus (PC, FCI, DirectLiNGAM, GRaSP), structurally distinct from the five-method consensus used on the lighter cohorts. The headline patient-stratified joint-causal-inference recall = 1.000 corner is robust to GES’s absence by construction of the union consensus (the Cross-context causal network section), but direction-accuracy and forbidden metrics on those cohorts could in principle shift if GES were included.
- **RESIT direction audit leaves a substantial fraction ambiguous.** Five of twelve evaluable directed edges in the universal headline are RESIT-ambiguous under the additive-noise model. The headline direction-accuracy numbers therefore reflect consensus-vote orientation, not independent residual-test confirmation; the RESIT annotations are reported as a per-edge confidence signal rather than as a correction layer.
- **Curated ground-truth coverage is incomplete.** The curated directed ground-truth set comprises 37 edges across the three contexts (normal, COVID-19, tumour), of which only 8 are evaluable on the 12-program joint-causal-inference pools; reporting evaluable recall rather than absolute recall partially controls for non-uniform program-set coverage but does not eliminate this limitation.
- **Algorithmic and program-aggregation assumptions.** The pipeline assumes causal sufficiency for PC, GES, GRaSP, and DirectLiNGAM (no unmeasured confounders); FCI partially relaxes this assumption by detecting potential latent confounders, but its output is consumed only as a binary skeleton vote in the consensus (the full partial-ancestral-graph markers are exported as a side artefact for downstream interrogation). The JASMINE scoring step additionally assumes program activity can be summarised by rank-aggregated gene-set expression within a target cell type; programs whose biological signal is carried by within-cell-type expression heterogeneity (for example, bistable switches) may be under-resolved by this aggregation.
- **Single tissue, single platform.** The 611-patient lung cohort is sourced exclusively from the CZ CELLxGENE Discover platform^24^. Cross-tissue and cross-platform replication is left to future work.
- **Observational causal claims.** All inferred edges are observational; the framework is hypothesis-generating and every reported edge — particularly the novel efferocytosis-driven CD4-activation edge and the cohort-discriminative-program arrows in Table 5b — requires experimental perturbation validation before any mechanistic claim is made.

## Conclusion

Cell-CLIP demonstrates that a mask-aware, multi-algorithm, joint-causal-inference-pooled causal-discovery framework can extract directed, mechanistically interpretable cell-program networks from patient-level single-cell atlases at evaluable recall = 1.000 with high direction accuracy and zero forbidden-orientation violations. The patient-stratified joint-causal-inference cross-context graph is the framework’s primary biological finding and the only configuration in the comprehensive ablation series to simultaneously reach all three optimal Pareto corners. We further identify a previously unreported failure mode of single-cohort causal inference on disease-extended pooled cohorts — a within-cohort mixing-induced direction collapse — and show empirically that joint-causal-inference pooling on patient-stratified cohorts dissolves it. By reporting per-cohort Pareto fronts rather than single best configurations, by documenting which cohorts are sample-size-limited and reporting them honestly, and by separating cross-context (patient-stratified joint-causal-inference) from single-cohort (universal majority, lung cancer patient-stratified union) reporting, Cell-CLIP advances computational tools for cell-cell communication toward principled, reproducible directional inference suitable for downstream perturbation design and in vitro / in vivo validation.

## Author Contributions

J.Z. conceived the study and designed experiments. J.Z. and M.K.F.A. conceived the software. M.K.F.A. implemented the software and performed experiments. J.Z. and X.L. wrote the manuscript with input from M.K.F.A.

## Competing Interests

Authors J.Z. and X.L. are Novartis employees.

## Data Availability

Code, sweep logs, curated ground-truth, and per-cohort detailed validation outputs available at the official publication.

## Acknowledgements

The authors thank Raf Luna, Joel Wagner, and Shiva Malek for their feedback, encouragement, and support during M.K.F.A.’s summer internship at Novartis Biological Research.

## Supplement

**Supplementary Figure S1.**
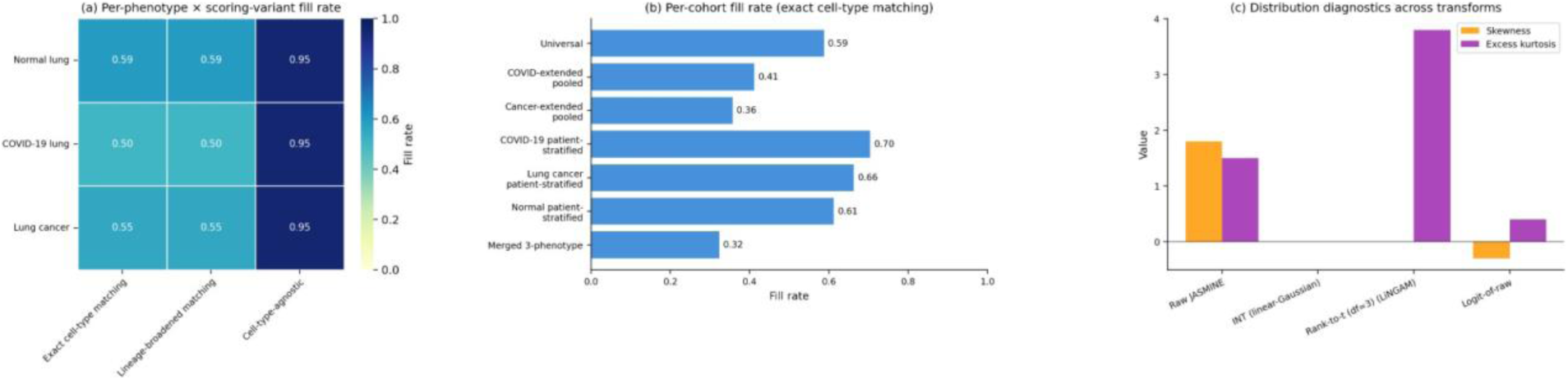
Mask-aware JASMINE diagnostics. **(a)** Per-phenotype × per-variant fill-rate matrix, showing how exact and lineage-broadened cell-type matching produce identical rates while the diagnostic-only cell-type-agnostic aggregation is substantially denser. **(b)** Per-cohort fill rates after cohort-filtering and patient-stratification. **(c)** Distribution diagnostics on the universal cross-phenotype matrix across four transforms (raw, INT, t (df = 3), logit), confirming INT yields skewness ≈ 0 and excess kurtosis ≈ 0 (correct input for Fisher’s-Z) while the t (df = 3) transform is symmetric with kurtosis ≈ 3.8 (correct input for DirectLiNGAM).

**Supplementary Figure S2.**
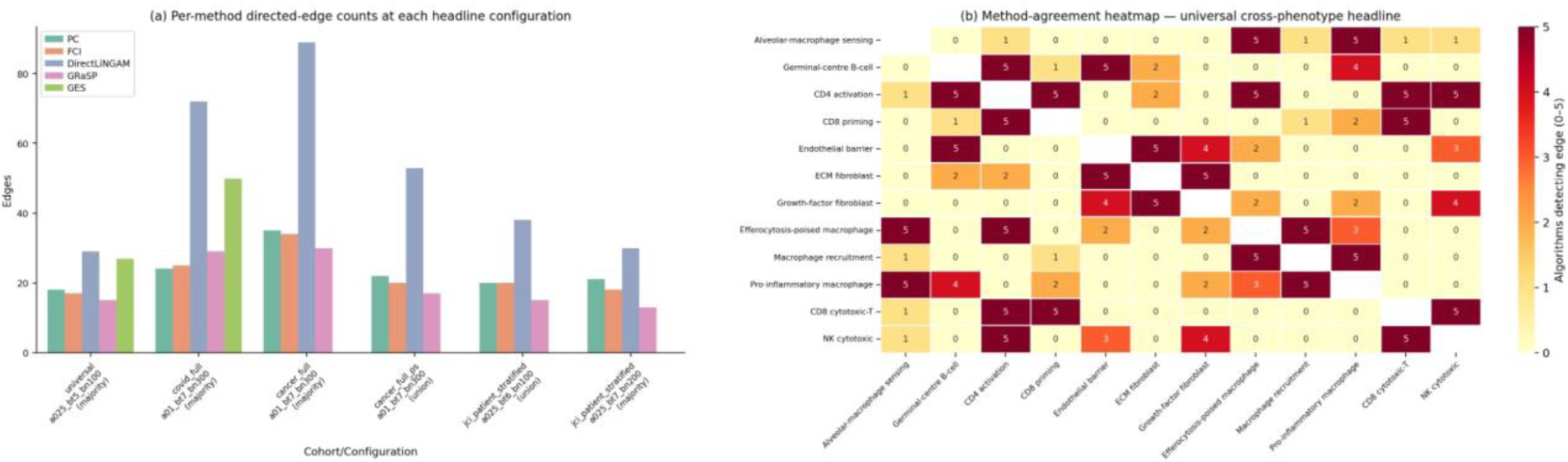
**(a)** Per-method directed-edge counts at each cohort’s headline hyperparameter grid point, broken down by PC, FCI, DirectLiNGAM, GRaSP, and GES. **(b)** Method-agreement heatmap on the universal cross-phenotype cohort (α = 0.025, bt = 0.5, bn = 100): each cell indicates the number of algorithms (0–5) supporting an edge between a pair of programs.

**Supplementary Figure S3.**
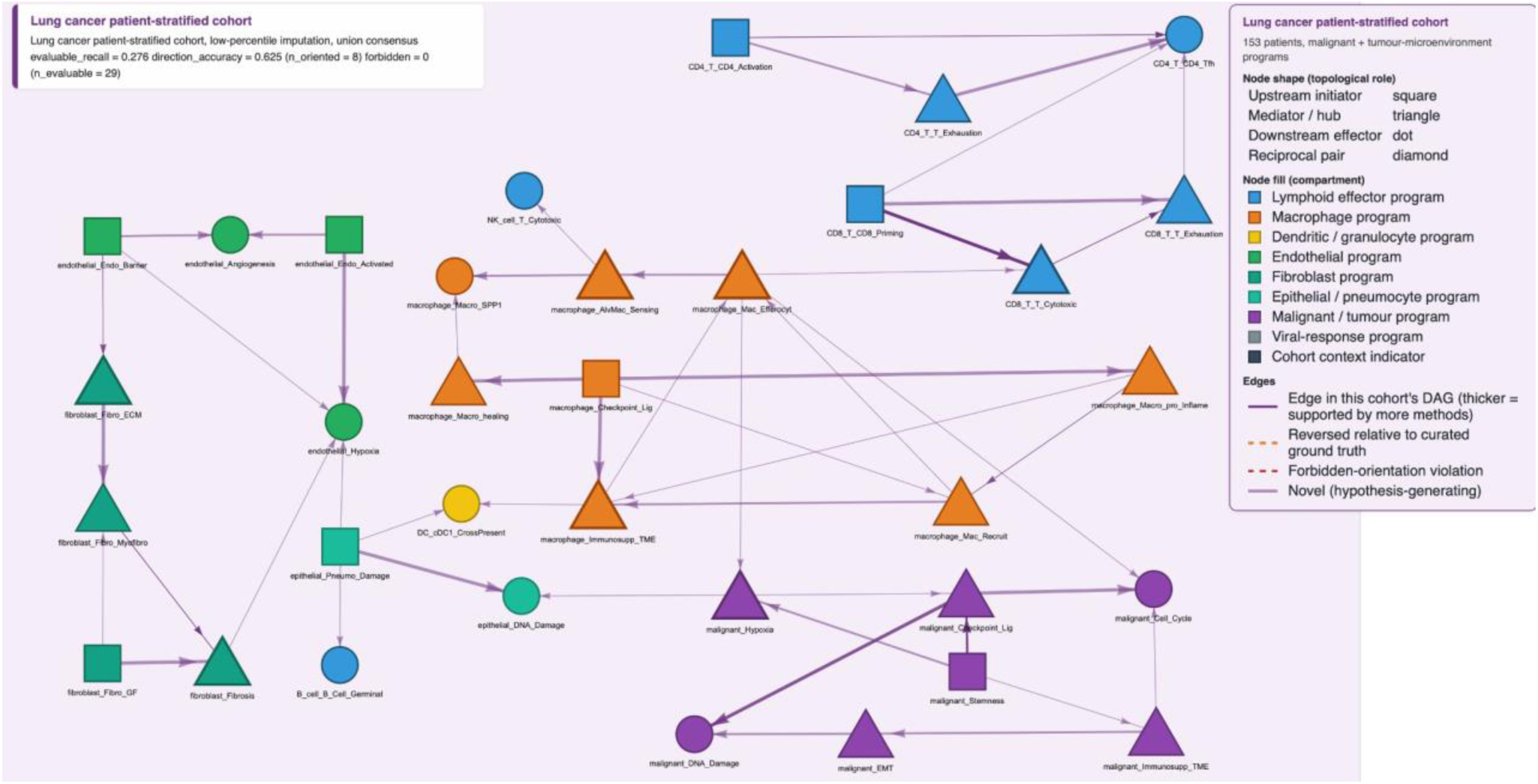
Lung cancer patient-stratified single-cohort union directed acyclic graph (hyperparameter setting α = 0.1, bootstrap threshold = 0.7, 300 bootstrap iterations; 49 edges; evaluable recall = 0.276, direction accuracy = 0.625, zero forbidden-orientation violations against the curated tumour-microenvironment ground-truth set). Node shape and fill encoding follow Figures 3 and 4. Edge colour is the lung cancer cohort accent (royal purple) for confirmed and novel orientations, dashed orange for reversed orientations. The graph contains malignant programs (purple fill) that are absent from the universal program set, including the stemness, checkpoint-ligand, hypoxia, epithelial-mesenchymal-transition, cell-cycle, DNA-damage-response, and immunosuppressive tumour-microenvironment programs.

**Supplementary Figure S4.**
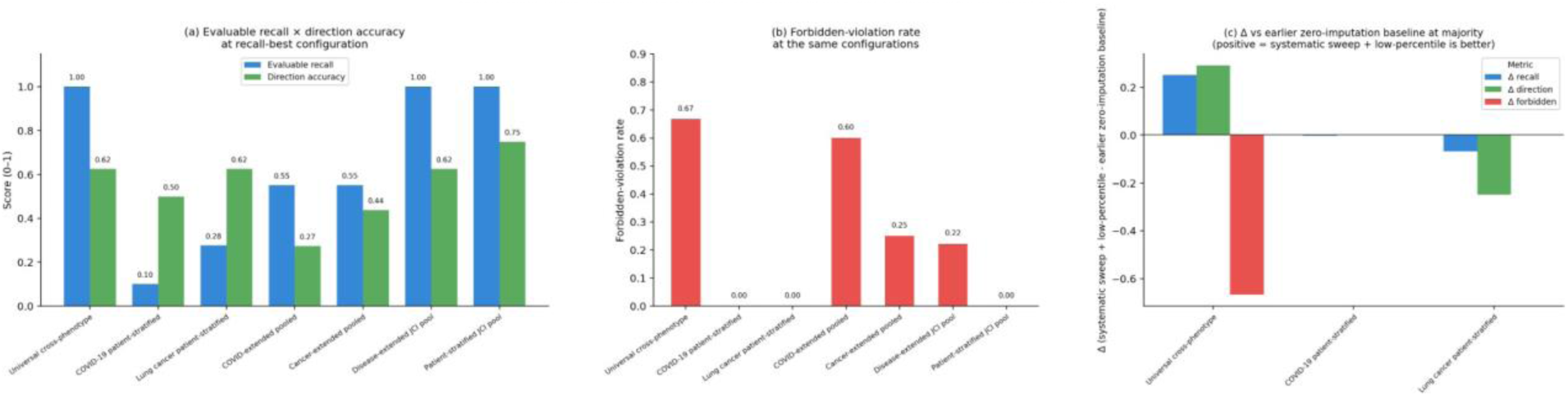
Validation metrics across reporting cohorts and consensus types. **(a)** Evaluable recall × direction accuracy at the recall-maximising configuration for each cohort. **(b)** Forbidden-violation rate at the same configurations. **(c)** Difference (Δ) versus the earlier zero-imputation baseline configuration at majority consensus, for the three cohorts that the earlier baseline covered (universal cross-phenotype, COVID-19 patient-stratified, lung cancer patient-stratified).

**Supplementary Figure S5.**
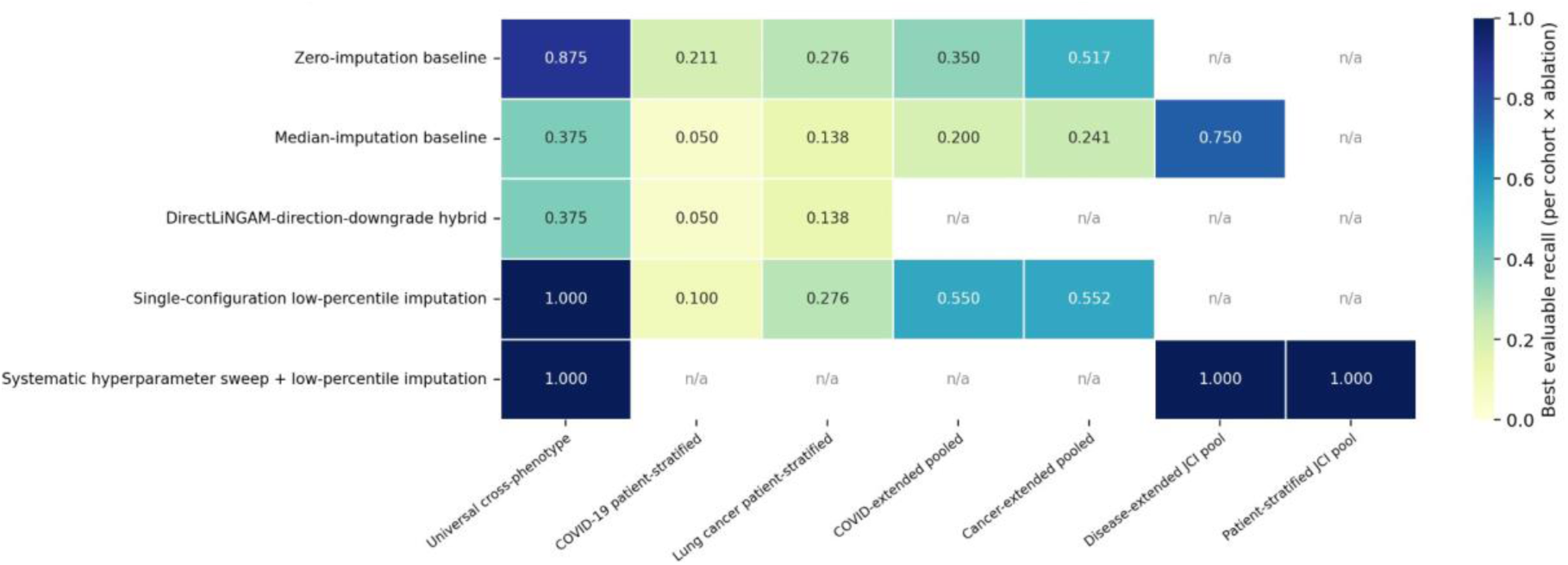
Ablation evaluable-recall heatmap. Rows: zero-imputation baseline, median-imputation baseline, DirectLiNGAM-direction-downgrade hybrid, single-configuration low-percentile imputation, systematic hyperparameter sweep with low-percentile imputation (variants defined in the Sensitivity analysis section). Columns: universal cross-phenotype cohort, COVID-19 patient-stratified cohort, lung cancer patient-stratified cohort, COVID-extended pooled cohort, cancer-extended pooled cohort, disease-extended cross-cohort JCI pool, patient-stratified cross-cohort JCI pool. Cell colour: best evaluable recall under each (row, column) cell.

## Notes

### Competing Interest Statement

Authors J.Z. and X.L. are affiliated with Novartis are employees.

### Summary of Updates

(i) updated abstract; (ii) text corrections and clarifications throughout the manuscript; (iii) added new Wilson confidence interval metrics; (iv) updated figure labels; and (v) fixed PDF formatting for improved readability and submission compliance.

